# Age and diet shape the genetic architecture of body weight in Diversity Outbred mice

**DOI:** 10.1101/2020.11.04.364398

**Authors:** Kevin M. Wright, Andrew Deighan, Andrea Di Francesco, Adam Freund, Vladimir Jojic, Gary Churchill, Anil Raj

## Abstract

Understanding how genetic variation shapes an age-dependent complex trait relies on accurate quantification of both the additive genetic effects and genotype-environment interaction effects in an age-dependent manner. We used a generalization of the linear mixed model to quantify diet-dependent genetic contributions to body weight and growth rate measured from early development through adulthood of 960 Diversity Outbred female mice subjected to five dietary interventions. We observed that heritability of body weight remained substantially high (*h*^2^ ≈ 0.8) throughout adulthood under the 40% calorie restriction diet, while heritability, although still appreciably high, declined with age under all other dietary regimes. We identified 14 loci significantly associated with body weight in an age-dependent manner and 19 loci that contribute to body weight in an age- and diet-dependent manner. We found the effect of body weight alleles to be dynamic with respect to genomic background, age, and diet, identifying the scope of pleiotropy and several instances of allelic heterogeneity. In many cases, we fine-mapped these loci to narrow genomic intervals containing a few genes and impute putative functional variants from the genome sequence of the DO founders. Of the loci associated with body weight in a diet-dependent manner, many have been previously linked to neurological function and behavior in mice or humans. These results enable us to more fully understand the dynamics of the genetic architecture of body weight with age and in response to different dietary interventions, and to predict the effectiveness of dietary intervention on overall health in distinct genetic backgrounds.

## 1 Introduction

Quantifying the contributions of genetic and environmental factors to population variation in an age-dependent phenotype is critical to understanding how phenotypes change over time and in response to external perturbations. The identification of genetic loci that are associated with a complex trait in an age- and environment-dependent manner allows us to elucidate the dynamics and context-dependence of the genetic architecture of the trait and facilitates trait prediction. For health-related traits, these genetic loci may also facilitate greater understanding and prediction of age-related disease etiology, which is an important step to genetically or pharmacologically manipulating these traits to improve health.

Standard approaches used to identify genetic loci associated with quantitative traits can be confounded by non-additive genetic effects such as genotype-environment (GxE) and genotype-age (GxA) interactions [1,], contributing to the “missing heritability” of quantitative traits [2,]. The linear statistical models routinely used in genetic mapping analyses do not account for variation in population structure between environments and polygenicity in GxE interactions. Population structure can substantially increase the false-positive rate when testing for GxE associations [3,]. Furthermore, not accounting for polygenic GxE interactions has the potential to incorrectly estimate the heritability of quantitative traits in the context of specific environments [2,]. To address these limitations, recent efforts have generalized standard linear mixed models (LMMs) with multiple variance components that allow for polygenic GxE interactions and environment-dependent residual variation [2, 4, 3, 5,]. Moreover, these generalized LMMs substantially increase the power to discover genomic loci that are associated with phenotype in both an environment-independent and environment-dependent manner.

In this study, we used a generalized LMM to investigate the classic quantitative trait, body weight, in a large population of Diversity Outbred (DO) mice. Body weight was measured longitudinally from early development to late adulthood, before and after the imposition of dietary intervention at six months of age. We expect diet and age to be important factors affecting body weight and growth rate; however, it remains to be determined how these factors will interact with genetic variation to shape growth. Two early studies found significant genetic correlations for body weight and growth rate during the first 10 weeks of mouse development, which supported the hypothesis that growth rates during early and late development were affected by pleiotropic loci [6, 7,]. Subsequent experiments found that the heritability of body weight increased monotonically with age throughout development: from 29.3% to 76.1% between 1 and 10 weeks of age [6,], from 6% to 24% between 1 and 16 weeks of age ([8]), and from 9% to 32% between 5 and 13 weeks of age [9,]. The heritability of growth rate also varied with age, but exhibited a peak of 24% at 3 weeks of age and then declined to nearly 4% at 16 weeks of age [8,]. The strength of association and effect size of QTLs for body weight and growth rate were specific to early or late ages and were inconsistent with the hypothesis that pleiotropic alleles affect animal size at early and late developmental stages [6, 8,]. While these results are well supported, their interpretation is somewhat limited because body weight measurements ceased at young ages and significant QTLs encompassed fairly large chromosomal regions. Given these limitations, we were motivated to ask two questions: Will fine-mapping to greater resolution reveal single genes which function at either early or late developmental stages, or reveal multiple genes in tight linkage with variable age-specific effects? How will the effect of these loci change at later ages and under different diets?

We expect the interaction of dietary interventions, such as caloric restriction or intermittent fasting, with genetics to greatly impact the body weight trajectories of mice. Researchers have observed genotype-dependent reductions in body weight in the 7 to 50 weeks after imposing a 40% caloric restriction, and variation in heritability of this trait with age from 42% to 54% [10,]. A second study subjected a large genetic-mapping population to dietary intervention and identified multiple loci with significant genetic and genotype-diet interaction effects on body weight at 2 to 6 months of age [11,]. These studies identified substantial diet- and age-dependent genetic variance for body weight in mice, similar to what has been found in humans [1, 12, 13,]. It, however, remains to be determined how the contribution of specific genetic loci to body weight changes in response to different dietary interventions and whether the effects observed in younger mice are indicative of the maintenance of body weight in adult mice.

In order to address these questions, we measured the body weight of multiple cohorts of genetically diverse mice from the DO population [14,] from 60 to 660 days of age (Figure 1A). At 180 days of age, we randomized mice by body weight and assigned each mouse to one of five dietary regimes – Ad libitum (AL), 20% and 40% daily calorie restriction (CR), and 1 or 2 day per week intermittent fasting (IF). Longitudinal measurements of body weight in these DO mice allowed us to discover the age-dependent genetic determinants of body weight and growth rate, in the context of different dietary interventions.

**Figure 1:**
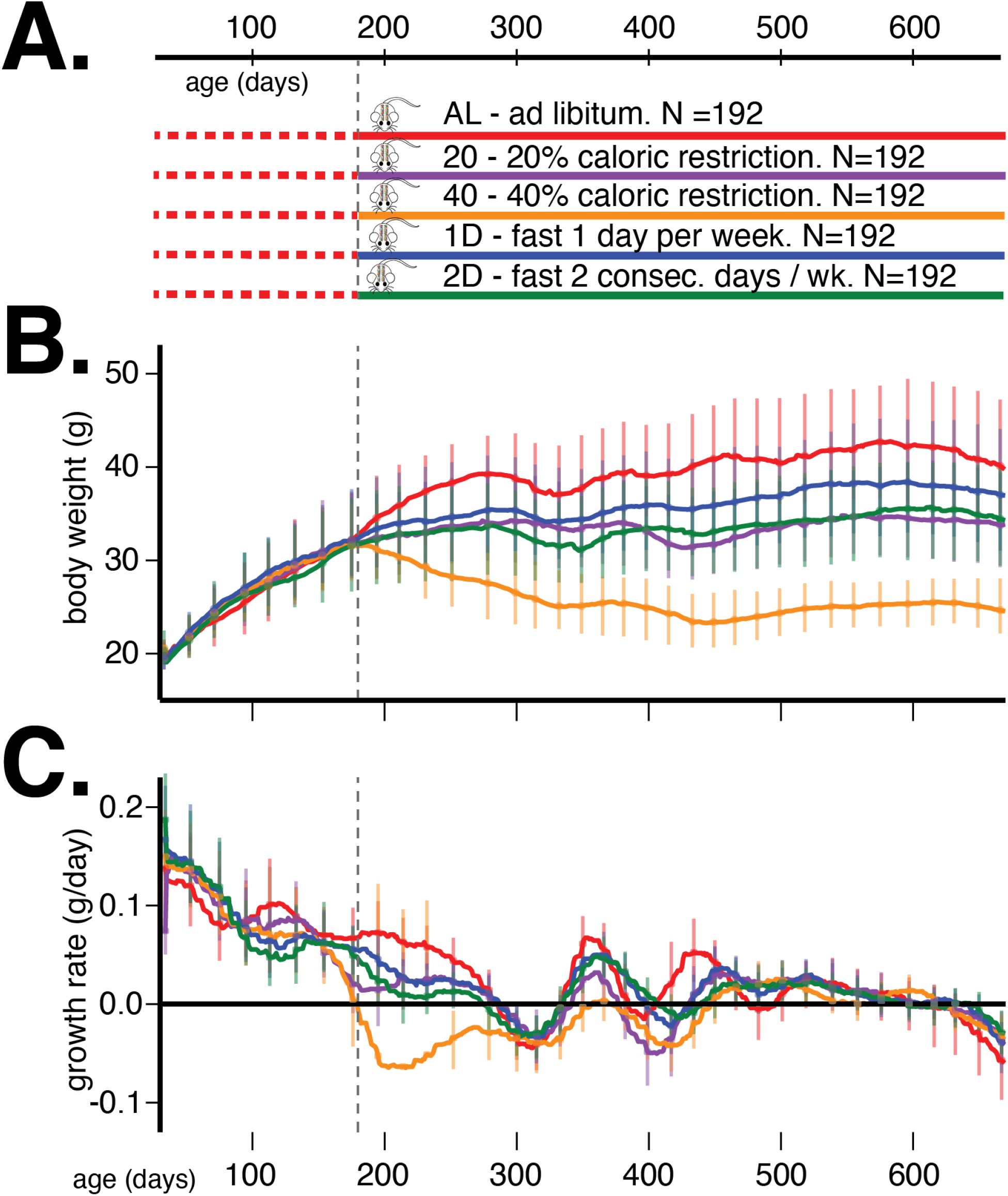
(A) Outline of study design. (B) Median (inter-quartile range) body weight in grams and (C) median (inter-quartile range) growth rate in grams per day for five dietary treatments from 60 to 660 days of age. Vertical grey dotted line denotes the onset of dietary intervention at 180 days of age.

In the following sections, we first describe the study design and collection of the genetic and body weight measurements. Second, we specify the Gene-Environment Mixed Model (GxEMM) [5,], the generalized LMM we use to quantify total and diet-dependent heritabilities of body weight and growth rate from 60 to 660 days of age. We next use this model to identify genetic loci having additive or genotype-diet interaction effects on body weight. We fine-map candidate loci and determine the scope of pleiotropy for age and diet specific effects. We find many, but not all, loci are associated with body weight in a narrow age range and localize to small genomic regions, in some cases to single genes. We utilize the full genome sequence of the DO founders and external chromatin accessibility data to further narrow the genomic regions to a small number of candidate variants at each locus. Interestingly, many diet-specific body weight loci localize the genes implicated in neurological function and behavior in mice or humans.

## 2 Study Design and Measurements

The Diversity Outbred (DO) house mouse (*Mus musculus*) population was derived from eight inbred founder strains and is maintained at Jackson Labs as an outbred heterozygous population [14,]. This study contains 960 female DO mice, sampled at generations: 22 – 24 and 26 – 28. There were two cohorts per generation for a total of 12 cohorts and 80 animals per cohort. Enrollment occurred in successive quarterly waves starting in March 2016 and continuing through November 2017.

A single female mouse per litter was enrolled into the study after wean age (3 weeks old), so that no mice in the study were siblings and maximum genetic diversity was achieved. Mice were housed in pressurized, individually ventilated cages at a density of eight animals per cage (cage assignments were random). Mice were subject to a 12 hr:12 hr light:dark cycle beginning at 0600 hrs. Animals exit the study upon death. All animal procedures were approved by the Animal Care and Use Committee at The Jackson Laboratory.

From enrollment until six months of age, all mice were on an Ad Libitum diet of standard rodent chow 5KOG from LabDiet. At six months of age, each cage of eight animals was randomly assigned to one of five dietary treatments, with each cohort equally split between the five groups (N=192/group): AD Libitum (AL), 20% caloric restriction (20), 40% caloric restriction (40), one day per week fast, (1D) and two days per week fast (2D) (see Figure 1A). In a previous internal study at the Jackson Laboratory, the average food consumption of female DO mice was estimated to be 3.43g/day. Based on this observation, mice on 20 diet were given 2.75g/mouse/day and those on 40 diet were given 2.06g/mouse/day. Food was weighed out for an entire cage of 8. Observation of the animals indicated that the distribution of food was roughly equal among all mice in a cage across diet groups.

Mice on AL diet had unlimited food access; they were fed when the cage was changed once a week. In rare instances when the AL mice consumed all food before the end of the week, the grain was topped off mid week. Mice on 20% and 40% CR diets were fed daily. We gave them a triple feeding on Friday afternoon to last till Monday afternoon. As the number of these mice in each cage decreased over time, the amount of food given to each cage was adjusted to reflect the number of mice in that cage. Fasting was imposed weekly from Wednesday noon to Thursday noon for mice on 1D diet and Wednesday noon to Friday noon for mice on 2D diet. Mice on 1D and 2D diets have unlimited food access (similar to AL mice) on their non-fasting days.

### 2.1 Body weight measurements

Body weight was measured once every week for each mouse throughout its life. The body weight measurements for this analysis were collated on February 1, 2020 at which point 941 mice (98%) had measurements at 180 days, 890(93%) at 365 days, 813 (85%) at 550 days, and 719(75%) at 660 days. For these analyses, we included all body weight measures for each mouse up to 660 days of age. We smoothed out measurement noise, either due to errors in measurement or swaps in assigning measurements to mice, using an *ℓ*_1_ trend filtering algorithm [15,] which calculates a piece-wise linear trend line for body weight for each mouse over its measurement span. The degree of smoothing was learned by minimizing the error between the predicted fit and measurements at randomly held-out ages across all mice. In the rest of this paper, *body weight* and *growth rate* refer to the predicted fits from *ℓ*_1_ trend filtering.

We present the average trends in body weight and growth rate, stratified by dietary intervention, in Figure 1 panels B and C, respectively. The body weight and growth rate trends without *ℓ*_1_ trend filtering are presented in Supplemental Figure S1. The most prominent observation from these trends is that dietary intervention contributes the most to variation in body weight in this mouse population. After accounting for this source of variation, there remains substantial and different quantities of variation in body weight trends between the different dietary interventions, suggesting a plausible GxD interaction effect on body weight trajectories.

### 2.2 Genotype measurements

We collected tail clippings and extracted DNA from 954 animals (http://agingmice.jax.org/protocols). Samples were genotyped using the 143,259-probe GigaMUGA array from the Illumina Infinium II platform [16,] by NeoGen Corp. (genomics.neogen.com/). We evaluated genotype quality using the R package: qtl2 [17,]. We processed all raw genotype data with a corrected physical map of the GigaMUGA array probes (https://kbroman.org/MUGAarrays/muga_annotations.html). After filtering genetic markers for uniquely mapped probes, genotype quality and a 20% genotype missingness threshold, our dataset contained 110,807 markers.

We next examined the genotype quality of individual animals. We found seven pairs of animals with identical genotypes which suggested that one of each pair was mislabelled. We identified and removed a single mislabelled animal per pair by referencing the genetic data against coat color. Next, we removed a single sample with missingness in excess of 20%. The final quality assurance analysis found that all samples exhibited high consistency between tightly linked markers: log odds ratio error scores were less than 2.0 for all samples [18,]. The final set of genetic data consisted of 946 mice.

For each mouse, starting with its genotypes at the 110,807 markers and the genotypes of the 8 founder strains at the same markers, we inferred the founders-of-origin for each of the alleles at each marker using the R package: qtl2 [17,]. This allowed us to test directly for association between founder-of-origin and phenotype (rather than allele dosage and phenotype, as is commonly done in QTL mapping) at all genotyped markers. Using the founder-of-origin of consecutive typed markers and the genotypes of untyped variants in the founder strains, we then imputed the genotypes of all untyped variants (34.5 million) in all 946 mice. Targeted association testing at imputed variants allowed us to fine-map QTLs to a resolution of 1 − 10 genes.

## 3 Models and Methods

### 3.1 Motivating models for environment-dependent genetic architecture

Genome-wide QTL analyses in model organisms over the last decade have predominantly employed linear mixed models (e.g., EMMA [19,], FastLMM [20,], and GEMMA [21,]), expanding on the heuristic that samples sharing more of their genome have more correlated phenotypes than genetically independent samples. We found that the distributions of covariances in body weight, measured at 500 days of age, between animal pairs within the AL treatment were nearly indistinguishable when we partition pairs into high kinship (>0.2) and low kinship groups (Figure 2, no significant separation between solid and dashed red lines). However, animal pairs in the 40%CR treatment exhibit significantly lower covariance in body weight in the low kinship group compared to the high kinship group (Figure 2, significant separation between solid and dashed orange lines). This observation suggests that the genetic contribution to body weight is different in distinct dietary environments. This observation also motivates the use of recently developed generalized linear mixed models to conduct genome-wide QTL analysis because they more fully account for the environment-dependent genetic variances, reduce false positive rates and increase statistical power [2, 4, 3, 5,]).

**Figure 2:**
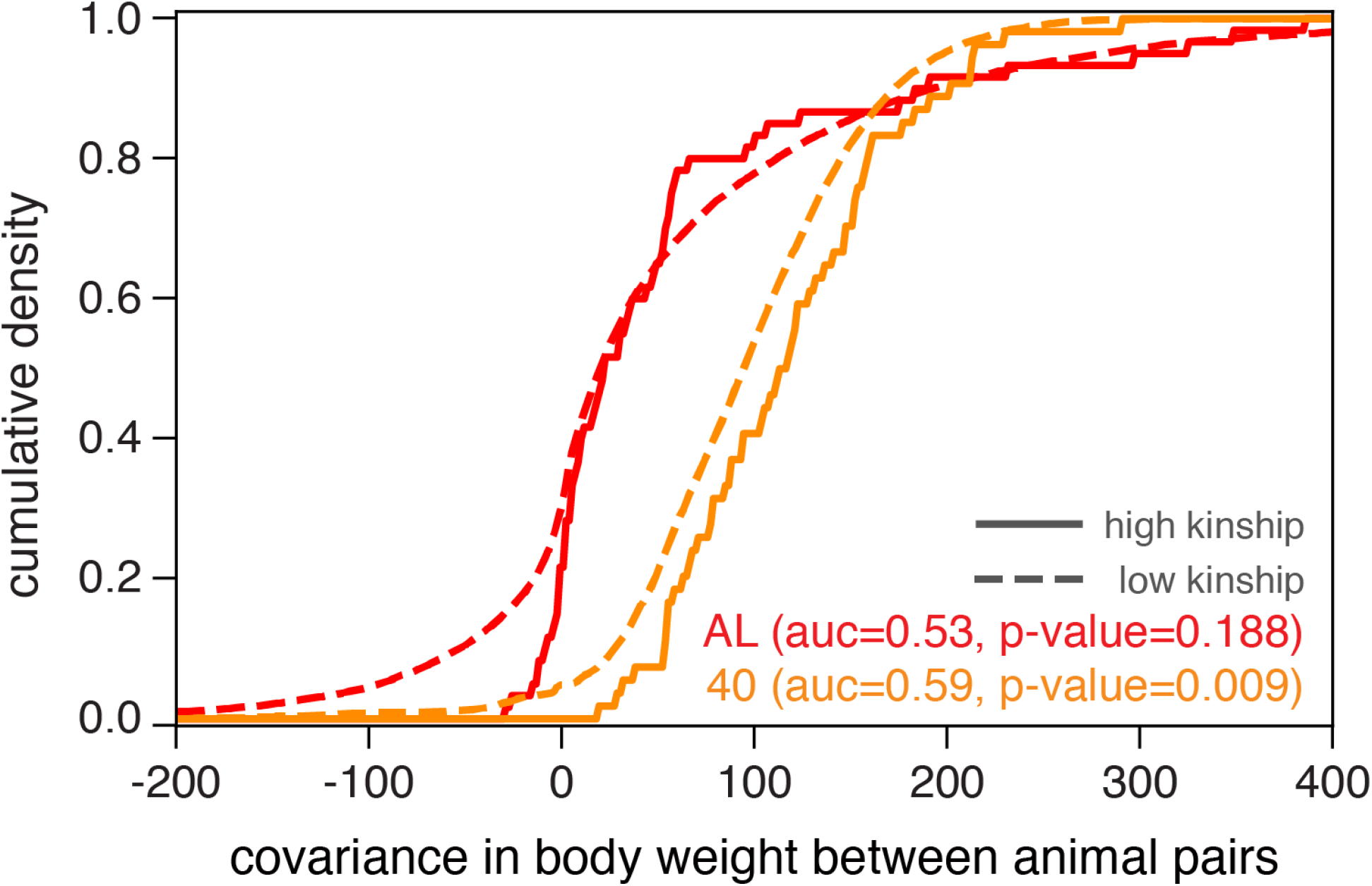
Phenotypic divergence between animal pairs is quantified by the covariance in body weight at 500 days of age. We plot the cumulative density of body weight covariance for all pairs of animals in the AL and 40% CR dietary treatments, partitioned into high kinship (> 0.2) or low kinship groups.

### 3.2 Overview of analyses

Starting with body weight measurements in 959 mice from 30-660 days of age, and founder-of-origin alleles inferred at 110,807 markers in 946 mice, we first quantified how the heritability of body weight and growth rate changes with age and between dietary contexts. We used the GxEMM model (described below in detail) to account for both additive environment-dependent fixed effects and polygenic gene-environment interactions. We considered two different types of environments: diet and generation. The five diet groups were assessed from 180 − 660 days and the twelve generations were assessed from 30 − 660 days. Next, we performed genome-wide QTL mapping for body weight at each age independently, testing for association between body weight and the inferred founder-of-origin at each genotyped marker. For ages 180 − 660 days, we additionally tested for association between body weight and the interaction of diet and founder-of-origin at each marker. We computed p-values using a sequential permutation procedure [22, 23,] at each variant for each of the additive and interaction tests and used these to assign significance [24,]. Finally, for each significant locus, we performed fine-mapping to identify the putative causal variants and founder alleles driving body weight, and underlying functional elements (genes and regulatory elements) to ascertain the possible mechanisms by which these variants act.

### 3.3 Polygenic models for gene x environment interactions

GxEMM [5,] is a generalization of the standard linear mixed model that allows for polygenic GxE effects. Under this model, the phenotype is written in terms of genetic effects as follows:

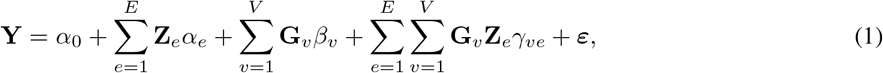

where **Y** ∈ ℝ^*N*^ is the vector of phenotypes over *N* samples, **G**_*v*_, *v* = 1, …, *V* are genotypes of *V* bi-allelic single nucleotide polymorphisms (SNPs), **Z**_*e*_, *e* = 1, …, *E* are binary vectors over *E* environments, and ***ε*** denotes the residual vector.

In our application, **Y** is the vector of *ℓ*_1_ trend filtered body weights at a specific age; we do not standardize the body weights so that the estimated effects are interpretable and comparable across ages. When testing for association with the founder-of-origin of markers, **G**_*nv*_ ∈ [0, 1]^8^; ||**G**_*nv*_||_1_ = 1 is a vector denoting the probability that the two alleles of the marker came from each of the eight founder lines from which the DO population is derived. ||·||_1_ denotes the *L*1-norm of a vector. Alternatively, when testing for association with the allele dosage of a variant, **G**_*nv*_ ∈ [0, 2] is the expected allele count at variant *v*. Finally, **Z**_*ne*_ = 1 denotes that sample *n* is subject to environment *e*. Prior to dietary intervention, the environments in our model are the 12 generations over which the DO samples span (i.e., *E* = 12). After dietary intervention, the environments further include the 5 diet groups (i.e., *E* = 17).

The effects of covariates, ***α***, are modeled as fixed while the genetic effects, ***β***, and genotype-environment effects, ***γ***, are modeled as random. Assuming heteroscedastic noise, 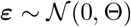, a normal prior on the random genetic effects, 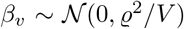, and a normal prior on the random GxE effects, 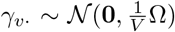, we get 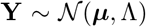, where ***μ*** = *α*_0_ + ∑_*e*_ **Z**_*e*_*α*_*e*_ and 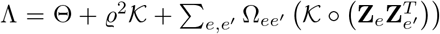. Θ is a diagonal matrix with entries specified as 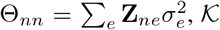 is the kinship matrix with entries defined as 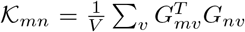, ϱ^2^ is the variance of environment-independent genetic effects, Ω ∈ ℝ^*E*×*E*^ is the variance-covariance matrix representing the co-variation in environment-dependent genetic effects between pairs of environments, and *A* ◦ *B* denotes the Hadamard product of matrices *A* and *B*. For simplicity, we constrain Ω to be a diagonal matrix in this study, limiting our ability to account for correlated genetic effects in pairs of environments. Note that each of the above parameters and data vectors in the model may be distinct at different ages.

### 3.4 Proportion of phenotypic variance explained by genetics

The proportion of total phenotypic variance explained by genetic effects is

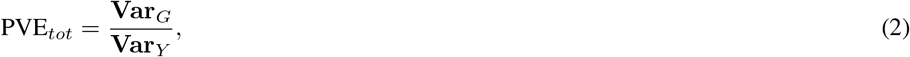

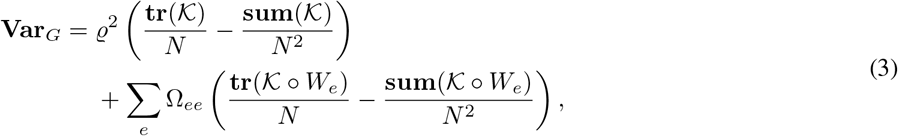

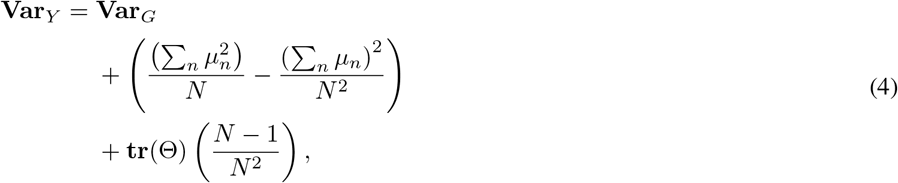

where 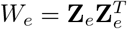, **tr**(·) denotes the trace of a matrix, and **sum**(·) denotes the sum of all elements of a matrix. The two terms in **Var**_*G*_ are genetic contributions to phenotypic variation that are shared across environments and specific to environments, respectively. The second and third terms in **Var**_*Y*_ are phenotypic variation explained by fixed effects and unexplained residual phenotypic variation, respectively.

Similarly, the environment-dependent PVE is

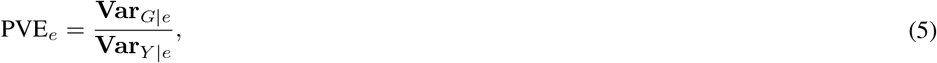

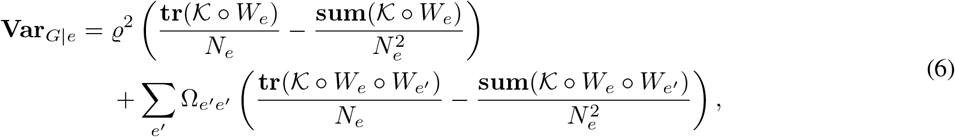

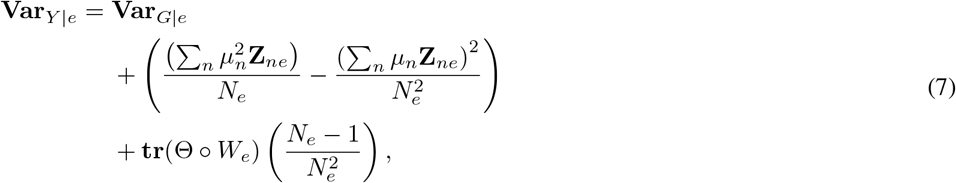

where *N*_*e*_ = ∑_*n*_ *Z*_*en*_ is the number of individuals in environment *e* (see Appendix A for more details). The proportion of phenotypic variance explained by genetic effects is equivalent to narrow-sense heritability, once variation due to additive effects of environment, batch, and other study design artifacts have been removed. In this work, we use the more general term, proportion of variance explained, to accommodate variation due to effects of diet and environment.

### 3.5 Genome-wide association mapping

#### 3.5.1 Additive genetic effects

To test for additive effect of a genetic variant on the phenotype, we include the focal variant among the fixed effects in the model while treating all other variants to have random effects.

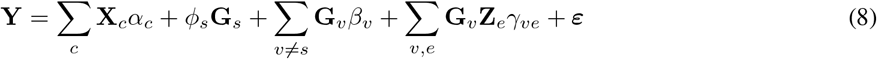

Applying the priors described above for *β*_*v*_, ***γ***_***v***_ and ***ε***, we can derive the corresponding mixed effects model is as follows:

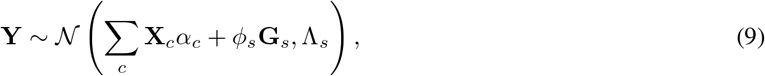

where 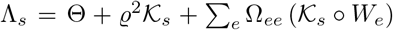 and 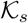 is the kinship matrix after excluding the entire chromosome containing the variant *s* (leave-one-chromosome-out or LOCO kinship). Leaving out the focal chromosome when computing kinship increases our power to detect associations at the focal variant [20,]. The test statistic is the log likelihood ratio Λ^*a*^(**Y**, **G**_*s*_) comparing the alternate model 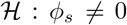 to the null model 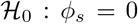. The log likelihood ratio is also referred to as log odds ratio or LOD through the rest of the paper.

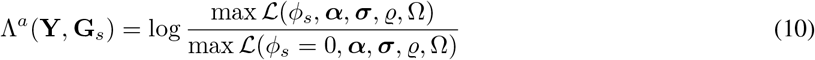

See Appendix B for details on computing the maximum likelihood estimates of the model parameters.

#### 3.5.2 Genotype-Environment effects

To test for effects of interaction between genotype and environment on the phenotype, we include a fixed effect for the focal variant and its interactions with the set of all environments of interest (*ε*) while treating all other variants to have random effects for their additive and interaction contributions,

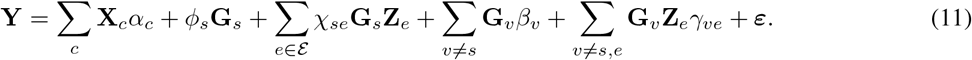

The corresponding mixed effects model is

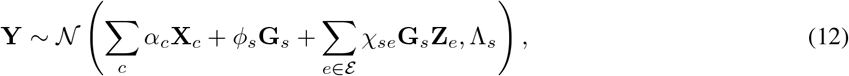

and the test statistic is

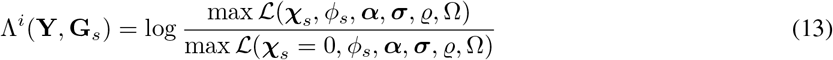

Note that this statistic tests for the presence of interaction effects between the focal variant and any of the environments of interest.

## 4 Simulations

To evaluate the accuracy of GxEMM and compare it against standard linear mixed models like EMMA ([19]), we simulated phenotypic variation under a broad range of values for PVE in each of two environments. For all simulations, we fixed the total sample size to *N* = 946 and used the observed kinship matrix for the 946 DO mice in this study. For each simulation, the environment-specific genetic contribution PVE_*e*_, environment-specific noise 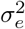, and the relative distribution of samples between environments are fixed. In order to explore how the two models behave under a wide range of conditions, we varied PVE_*e*_ ∈ {0.05, 0.10, …, 0.95}, 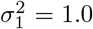, and 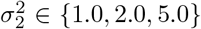, and the samples were assigned to environments either at a 1:1 ratio or a 4:1 ratio (i.e., a total of 114 parameter values). For each set of fixed parameter values, we run 100 replicate simulations. At the start of each simulation, we randomly assign samples to one of the two environments. Using equations 5, 6, and 7, we computed the variance component parameters by solving a set of linear equations. We then generated the vector of phenotypes from a multivariate normal distribution with zero mean and covariance structure dependent upon these variance component parameters (as per the generative model above). Using the simulated phenotypes, we estimated PVE_*tot*_ using both the EMMA and GxEMM models and PVE_*e*_ using the GxEMM model.

We first evaluated the accuracy of PVE_*tot*_ estimated from the two models and found that the GxEMM model estimated total PVE with little bias and lower variance compared to EMMA in nearly all simulations across the suite of parameter combinations that we investigated (Figures 3A and 3B). In particular, we found that EMMA substantially underestimated the total PVE in comparison to GxEMM when samples were equally distributed between environments and there was substantial difference in PVE or noise between environments. This is because PVE_*e*_ will have a greater impact on PVE_*tot*_ when both environments are more equally represented in the study sample.

**Figure 3:**
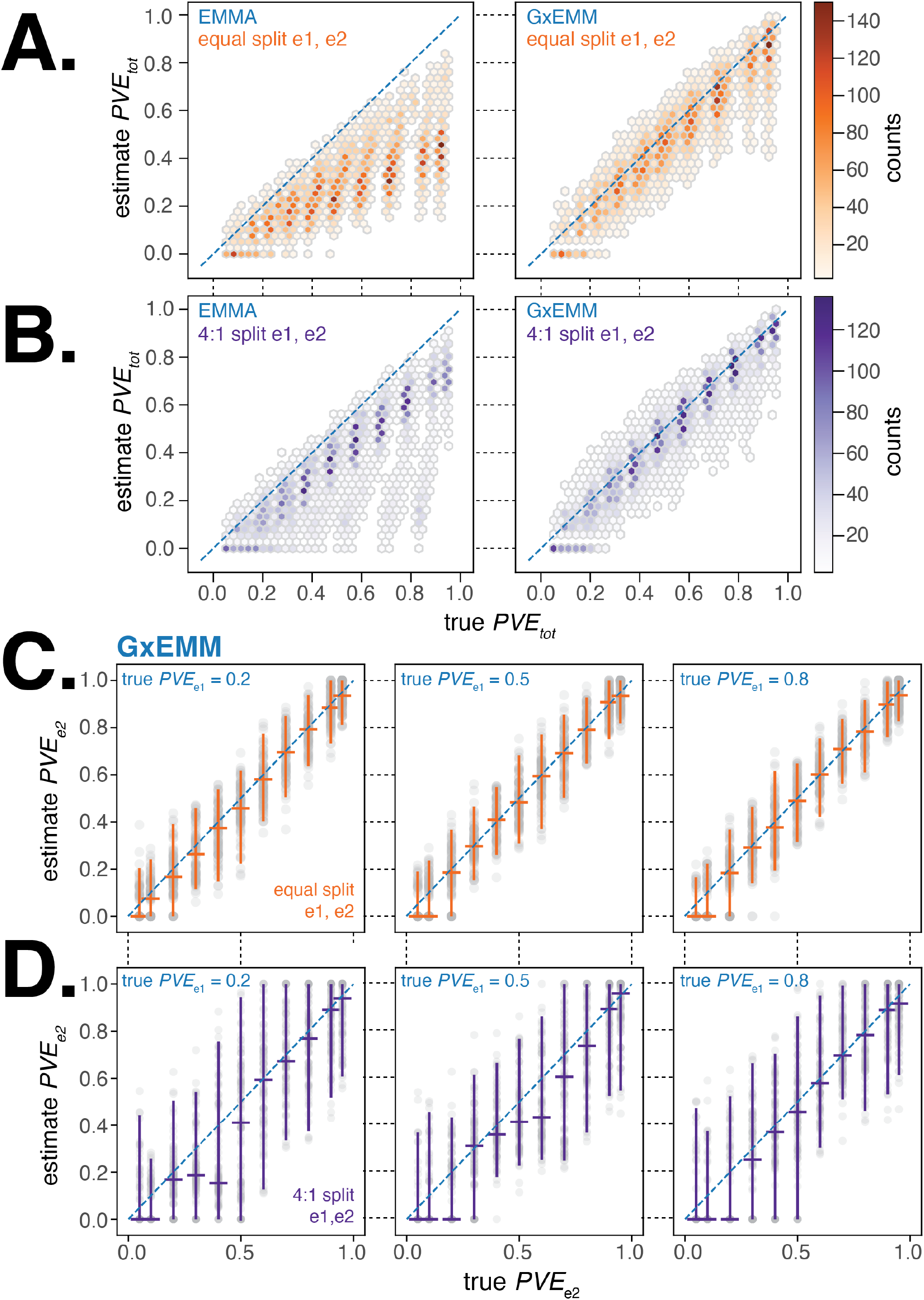
Evaluating GxEMM and EMMA using simulated datasets. (A). Comparison of true PVE_*tot*_ to that estimated from EMMA (left panel) and GxEMM (right panel) models. Simulations were run with an equal number of samples each environment and 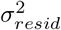 was fixed at 1.0. We ran for pairwise combination of PVE_*e*1_ and PVE_*e*2_ varied from 0.05 to 0.95, with fifty replicates per pair of values. (B). Same as (A) but with a 4:1 of samples between environments 1 and 2. (C). Plot of the true vs GxEMM estimate of PVE, the PVE specific to environment 2.

Next, we examined the sensitivity and specificity of the GxEMM model to quantify environment-specific genetic contribution to phenotypic variance. We found that when samples were equally distributed between environments, GxEMM accurately estimates PVE_*e*_ under a wide range of parameter values (Figure 3C). We observe an exception to this general result wherein we underestimate PVE_*e*_ for the environment with lower PVE when there is a large difference in PVE between the two environments. In contrast, when there is a skew in the number of samples between environments, we tend to underestimate low PVE values for the environment with fewer samples (Figure 3D). We observe similar patterns of bias when the two environments have comparable PVE values but different amounts of environment-specific noise.

Overall, the GxEMM model outperforms the EMMA model in estimating total PVE and shows little bias in estimating environment-specific PVE across a broad range of scenarios relevant to our study.

## 5 Results and Discussion

### 5.1 PVE of body weight across age and diet

First, we quantified the overall contribution of genetics to variation in body weight and, importantly, how this contribution changed with age. We applied both EMMA and GxEMM to body weight estimated every 10 days. Since the mice from each generation cohort were measured at the same time every week, we used generation as a proxy for the shared environment that mice are exposed to as part of the study design. We accounted for generation-specific fixed effects (***α*** in Equation 1) in both models and genotype-generation random effects (***γ***) in GxEMM. For ages after dietary intervention (≥ 180 days), we accounted for diet-specific fixed effects in both models and genotype-diet random effects in GxEMM. We estimated the variance components in the model at each age independently and computed the total and diet-dependent PVE using Equations 2 and 5.

In Figure 4A, we observed that the PVE of body weight steadily increased during development and up to 180 days of age, when dietary intervention was imposed. The GxEMM model estimates a higher PVE than the EMMA model during this age interval because the former model specifically accounts for polygenic genotype-generation effects. Following dietary intervention, PVE decreased in four of the five dietary groups; the one exception was the 40% CR group which maintained a high PVE (PVE_40_ ≈ 0.8) (Figure 4A) and low total phenotypic variance (Supplemental Figure S3A, top right panel) from 180 − 660 days of age. In contrast, the Ad-lib group had the lowest PVE and greatest total phenotypic variance in the same age range. Notably, *ℓ*_1_ trend filtering of the raw measurements proved useful in quantifying smoothly varying trends in the PVE of body weight and growth rate, and improving our estimates of the PVE of growth rate by reducing the effect of measurement noise (Supplemental Figure S2). Moreover, these results were robust to variation in the genetic data used to calculate the kinship matrix and to survival bias at 660 days of age.

**Figure 4:**
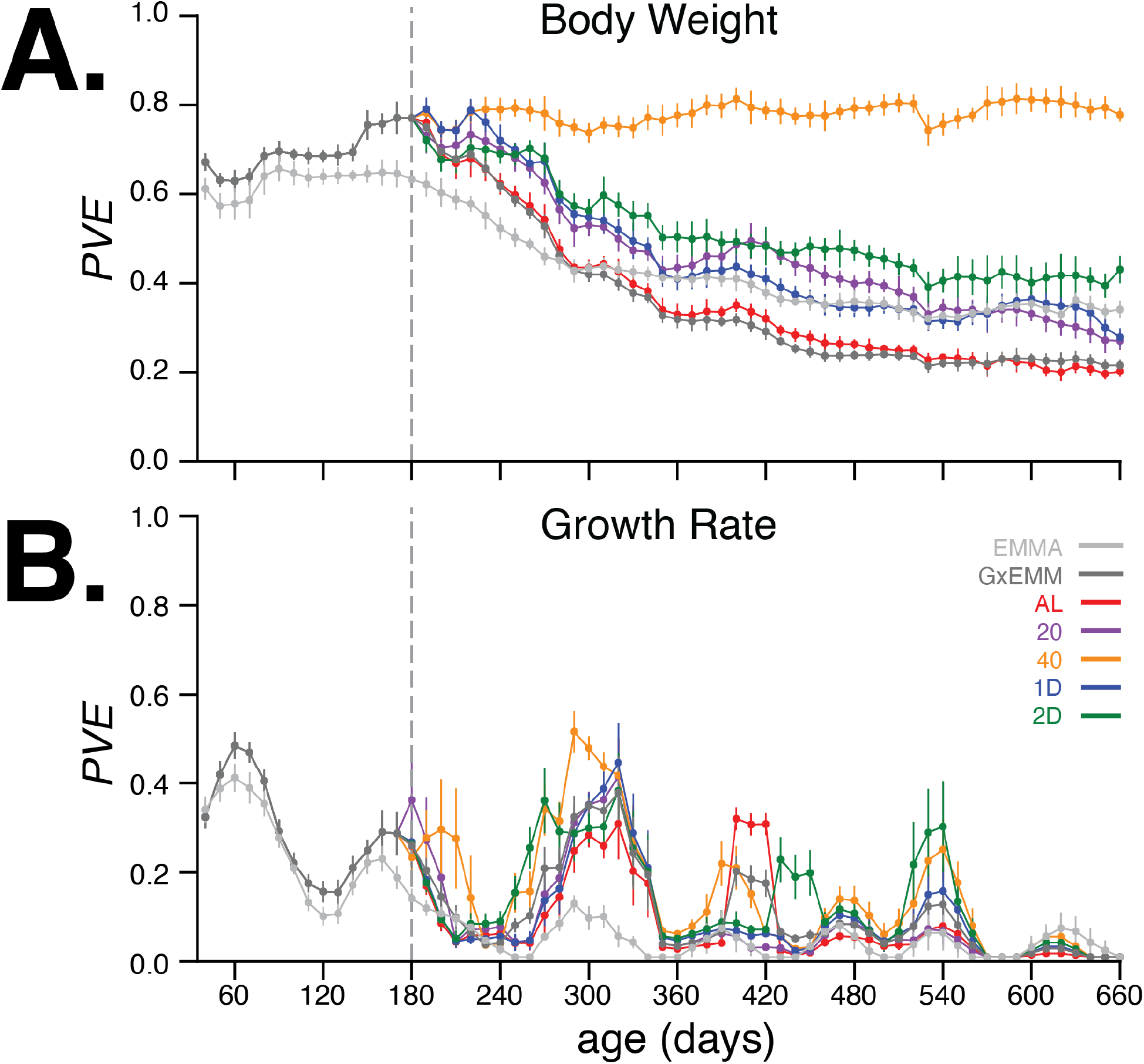
(A) Body weight PVE values for 30 − 660 days of age. Total PVE estimates are derived from the EMMA (light grey) and GxEMM (dark grey) models. Diet-dependent PVE values are derived from the GxEMM model. Dotted vertical line at 180 days depicts the time at which all animals were switched to their assigned diets. (B) Growth rate PVE; details the same as A.

The kinship matrix used for estimating these PVE values was computed based on the founder-of-origin of marker variants [25,]. When using kinship estimated using bi-allelic marker genotypes (as is commonly done in genome-wide association studies), we observed largely similar trends in PVE; however, differences in PVE between diets after 400 days were harder to discern due to much larger standard errors for the estimates (Supplemental Figure S3A, left panels). To test for bias or calibration errors in our PVE estimates, we randomly permuted the body weight trends between mice in the same diet group and re-calculated the total and diet-dependent PVE values. Consistent with our expectations, PVE dropped to nearly zero for the permuted dataset (Supplemental Figure S3B, left panel), indicating that the PVE estimates are well-calibrated. Note that when using kinship computed from genotypes, the PVE in the permuted dataset does not drop to zero, suggesting that PVE estimated in this manner is not well-calibrated (Supplemental Figure S3B, right panel). Finally, to evaluate the contribution of survival bias to our estimates, we recomputed PVE at all ages after restricting the dataset to mice that were alive at 660 days. We observed PVE estimates largely similar to those computed from the full dataset, suggesting very little contribution of survival bias to our estimates (Supplemental Figure S3C).

Next, we quantified the age-dependent contribution of genetics to variation in growth rate, enabled by the dense temporal measurement of body weight. As before, we applied EMMA and GxEMM to growth rate estimated every 10 days. In Figure 4B, we observed that PVE of growth rate increases rapidly during early development, and then decreases to negligible values around 240 days of age. In contrast to body weight, PVE of growth rate is substantially lower at all ages, and there is little divergence in PVE across diet groups for most ages. Notably, the decrease and subsequent increase in growth rate PVE coincides with specific metabolic, hematologic and physiological phenotyping procedures that these mice underwent at specific ages as part of the study (Supplemental Figure S1). Due to lower values and greater variance in PVE of growth rate with age, we focus on body weight throughout the rest of the paper.

In summary, the 40% CR intervention produced the greatest reduction in average body weight and maintained a high PVE after dietary intervention. This is because the total genetic variance in body weight remained relatively high and the environmental variance remained relatively low throughout this interval. In contrast, body weight PVE steadily decreased with age in each of the four less restrictive diets, which was due to a steady increase in environmental variance and not a decrease in the total genetic variance of body weight (Supplemental Figure S3A). Even though the total genetic variance in body weight is nearly constant across diets from 180 − 660 days of age, the effect of specific variants may change with age.

### 5.2 Genomewide QTL analysis of body weight across age and diet

We sought to identify loci significantly associated with body weight in a diet-dependent and age-dependent manner. To this end, we tested for association between the inferred founder-of-origin of each typed variant and body weight at each age independently. We note that body weight measurements taken at different ages are not independent; i.e., we may detect a locus at a specific age if it has small effects on body weight acting over a long period of time or a large effect on body weight resulting in rapid bursts in growth. Thus, a locus identified as significant at a specific age indicates its cumulative contribution to body weight at that age.

We identified 29 loci significantly associated with body weight at any age in the additive or interaction models using a *p* ≤ 10^−4^ cutoff for the additive test and a weaker *p* ≤ 10^−3^ cut-off for the genotype-diet interaction test (Supplemental Figure S4). Using all bi-allelic variants imputed from the complete genome sequences of the eight DO founder strains [26,], we re-tested the genetic association for the 29 candidate loci with the additive and interaction models. Specifically, we tested for association between all imputed genotype variants and body weight, accounting for kinship estimated using founder-of-origin inferred at genotyped variants as before. We found that 24 loci remained significant in the fine mapping analysis: five loci unique to the additive model, ten loci unique to the interaction model, and nine loci significant in both models (Table 1). We identified body weight associations with age-dependent effects from early development to adulthood: four diet-independent loci were associated with body weight exclusively during development (ages 60 – 160 days) and three were associated exclusively during adulthood (ages 200 – 660 days); the remaining seven loci were associated during development prior to the imposition of dietary restriction at day 180 and continued to be associated into adulthood (Table 1). The majority of diet-dependent loci (12 of 19) had a detectable effect on body weight only 240 days after dietary intervention (ages 420 – 660 days; Table 1). For each candidate body weight locus, we sought to determine the likely set of causal variants and estimate the effect of the eight founder alleles in a diet- and age-dependent manner.

**Table 1:**
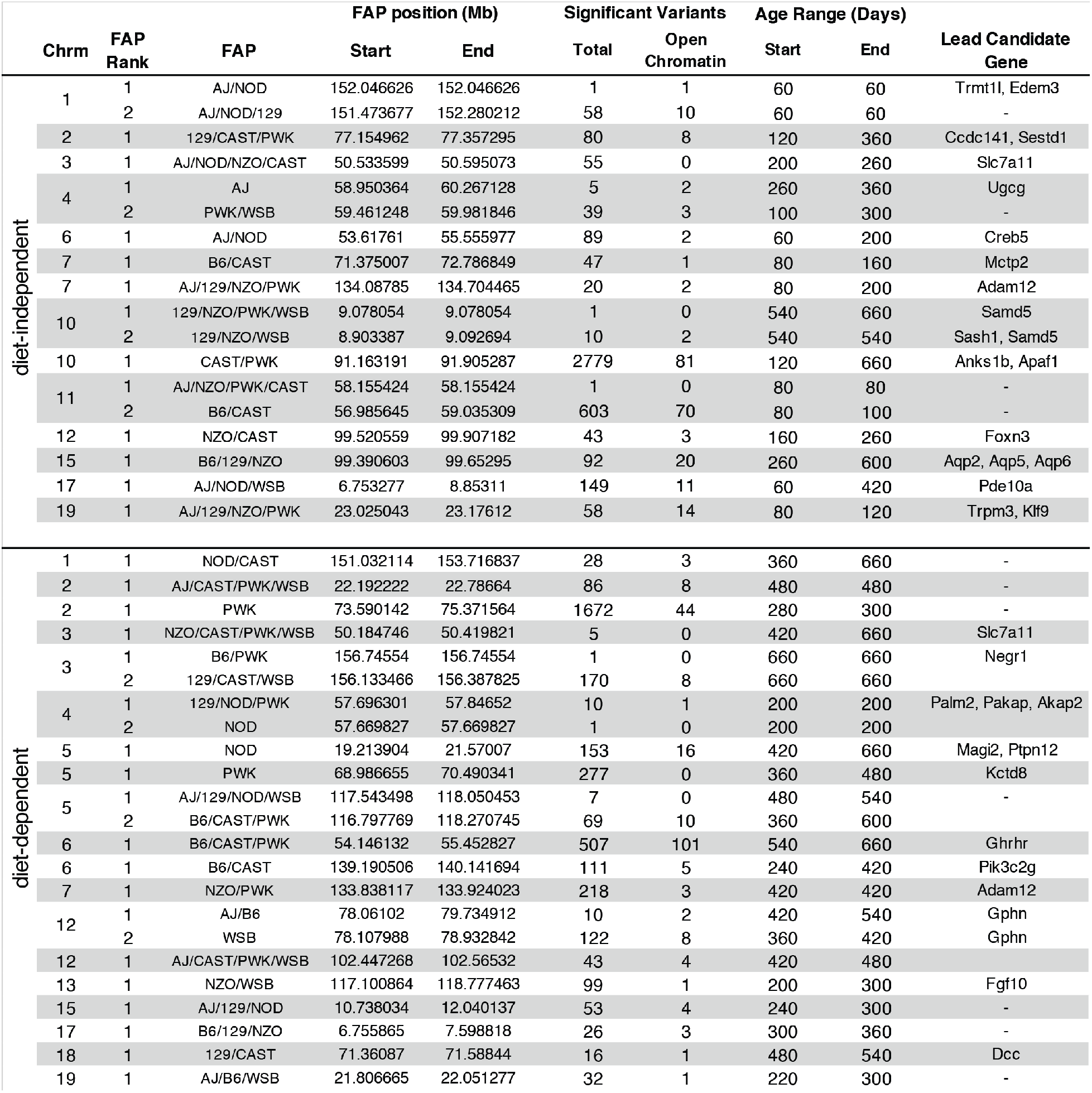
Significant body weight loci identified using the additive and interaction models. For each locus, we identified the founder allele pattern (FAP) of the variant with the strongest association at any age, the genomic location of these variants, number of significant variants that comprise this lead FAP, the number of these variants located within regulatory elements identified using chromatin accessibility measurements, and the ages at which at least one variant in the lead FAP is significantly associated with body weight. For loci in which the lead FAP is comprised of fewer than 10 variants, we also present results for the second FAP. We list candidate genes for loci where the FAP spans three or fewer genes.

In order to facilitate characterization and interpretation of the genetic associations at each locus, we sought to represent these associations in terms of the effects of founder haplotypes. First, we determined the founder-of-origin for each allele at every variant that was significant in at least one age. This allowed us to assign a founder allele pattern (FAP) to each significant variant in the locus. For example, if a variant with alleles A/G had allele A in founders AJ, NZO, and PWK and the allele G in the other 5 founders, then we assign A to be the minor allele of this variant and define the FAP of the variant to be AJ/NZO/PWK. Next, we grouped variants based on FAP and define the LOD score of the FAP group to be the largest LOD score among its constituent variants. (Note that, by definition, no variant can be a member of more than one FAP group.) By focusing on the FAP groups with the largest LOD scores, we significantly reduced the number of putative causal variants (Table 1, Supplemental File 1), while representing the age- and diet-dependent effects of these loci in terms of the effects of its top FAP groups. We further narrowed the number of candidates by intersecting the variants in top FAP groups with functional annotations (e.g., gene annotations, regulatory elements, tissue-specific regulatory activity, etc). For many loci, this procedure identified candidate regions containing one to three genes (Table 1) and we provide the full list of all genes within candidate regions in Supplemental File 2. In order to demonstrate the utility of this approach, we first examined a single locus on chromosome 6 strongly associated under the additive model.

We identified a diet-independent locus on chromosome 6 significantly associated with body weight during early development and nominally associated with body weight in certain dietary treatments at later ages (Figure 5A). One explanation for this result is a single pleiotropic allele affecting body weight at two distinct stages of life: early development and adulthood. Alternatively, this result could be explained by allelic heterogeneity [27,] – a single locus harboring multiple functional alleles each with distinct phenotypic effects. A third possibility is that this single genomic region contains multiple functional body weight loci that are only revealed with sufficient fine-mapping resolution. Fine-mapping this locus using the additive model, we identified the variant with the highest LOD score to be at 53.6 Mb. The minor allele at this lead variant was common to the AJ and NOD founders, while the remaining 6 founders possessed the alternate allele; this defined the lead diet-independent FAP at this locus to be AJ/NOD (Figure 5B). Separately, we fine-mapped this locus using the interaction model, identified the lead variant at 55.1 Mb, and determined the lead FAP to be B6/CAST/PWK (Figure 5C). These results are consistent with the hypothesis that at least two distinct body weight QTLs with functional alleles derived from different DO founders were responsible for the diet-dependent and diet-independent body weight associations at this locus.

**Figure 5:**
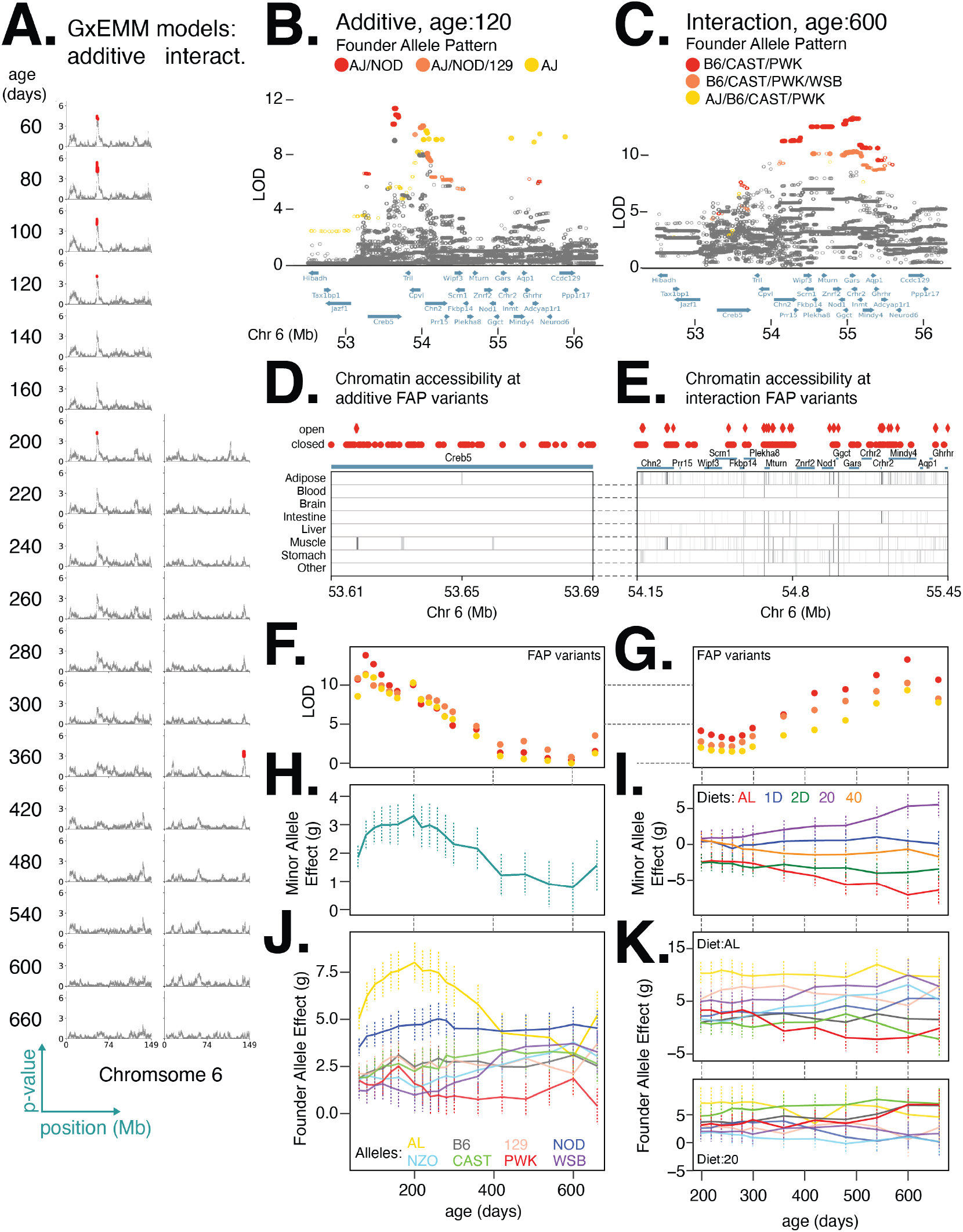
Distinct loci contribute to diet-independent and diet-dependent effects on body weight within a region on chromosome 6. (A) Manhattan plots of additive genetic associations and genotype-diet associations on chromosome 6 at multiple ages. (B) and (C) Fine-mapping loci associated with body weight, independent of diet at 80 days of age and dependent on diet at 600 days of age. Significant variants (solid circles). Colors denote variants with shared FAPs; rank 1, 2, and 3 by LOD score are colored red, orange, and yellow, respectively. (D) and (E) Significant variants, colored by their FAP group, along with the gene models (shown in green) and the tissue-specific activity of regulatory elements near these variants (shown in grey). Significant variants that lie within regulatory elements are highlighted as diamonds, and regulatory elements that contain a significant variant are highlighted in dark grey. (F) and (G) Log odds ratio of body weight association as a function of age for the lead variant from each FAP group. FAP colors are consistent with (B) and (C), respectively. (H) and (I) Estimated mean (se) effect on body weight (grams) of the minor allele for the lead imputed variant. For the diet dependent locus (I), effects are shown for each diet treatment. (J) Estimated mean (se) effect on body weight (grams) of each founder allele for the genotyped marker with the highest LOD score from the additive model. (K) Estimated effect (se) on body weight (grams) of each founder allele under the AL and 20% CR diets.

We hypothesized that the functional variant(s) responsible for the diet-independent and diet-dependent body weight associations at this locus are among the variants in the respective lead FAP groups because they exhibit the strongest statistical association and it is unlikely any additional variants are segregating in this genomic interval beyond those identified in the full genome sequences of the eight founder strains. For the diet-independent locus, at age 120, we identified 87 significantly associated variants; of these, 79 could be assigned to the lead FAP group and shared a similarly high LOD score (Figure 5B). All of these variants are SNPs located in the gene CREB5, 78 are intronic and one a synonymous exon variant. For the diet-dependent locus, at age 600 days, we identified 617 variants as significant; of these, 507 could be assigned to the lead FAP and shared a similarly high LOD score (Figure 5C). Two of the 507 variants were intergenic structural variants; of the remaining SNPs, 5 were non-coding exon variants, 167 were intronic, and the remainder were intergenic. Given that all candidate variants were non-coding, we next sought to determine whether they were located in regulatory elements across a number of tissues, identified as regions of open chromatin measured using ATAC-seq [28,] or DNase-seq [29,]. For the diet-independent and diet-dependent loci, we found 2 and 101 variants, respectively, that were located in regions of open chromatin (Figure 5D,E). Notably, both variants with diet-independent effects lay within the same muscle-specific regulatory element located within CREB5, suggesting that these variants likely affected body weight by regulating gene expression in muscle cells.

We found the relationship between model LOD score and age was similar for each of the three lead FAPs at the diet-independent and diet-dependent loci (Figure 5F, G). The minor allele of the lead variant at the diet-independent locus was associated with increased body weight at young ages (Figure 5H) whereas the minor allele for the lead diet-dependent variant had a positive effect on body weight under the 20% CR diet, a nearly neutral effect under the 40% CR, and a negative effect under the AL diet (Figure 5I). We next measured the effect of each founder allele at the lead diet-independent genotyped marker and, consistent with the lead FAP group for the diet-independent association, the AJ and NOD alleles had large positive effects at young ages (Figure 5J). For the diet-dependent association, B6, CAST, and PWK alleles were associated with decreased body weight in the AL diet and increase in body weight in the 20% CR diet (Figure 5K), consistent with their role in defining the lead diet-dependent FAP group. This example clearly demonstrates the insight gained by focusing on lead FAP variants to link specific founder alleles to variation in body weight and narrow the number of potential functional variants underlying body weight.

Next, we used this approach to examine a locus on chromosome 12 with diet-dependent effects on body weight from 300 to 420 days of age (Table 1; Supplemental Figures S5D). Upon fine-mapping the locus at 420 days of age, when the association was the strongest, we identified 77 significant variants partitioning into two distinct lead FAP groups, with similarly high LOD scores and centered at the same gene. The rank 1 FAP group contained variants with the minor allele specific to AJ and B6, whereas the rank 2 FAP group contained variants with the minor allele specific to WSB (Figure 6A). Of the 77 variants, the AJ/B6 FAP group contained 6 intergenic SNPs and 4 intronic SNPs spanning three genes: gephyrin (GPHN), Plekhh1, and RAD51b (Figure 6A). One of these ten variants is located in a regulatory element active specifically in adipose tissue. The remaining 67 significant variants all belonged to the WSB-specific FAP group; of these, 23 SNPs were intergenic and 44 were intronic and centered at the gephyrin gene. Four of these 67 variants were located in regulatory elements active in adipose tissue as well as other tissues relevant to metabolism (Figure 6B).

**Figure 6:**
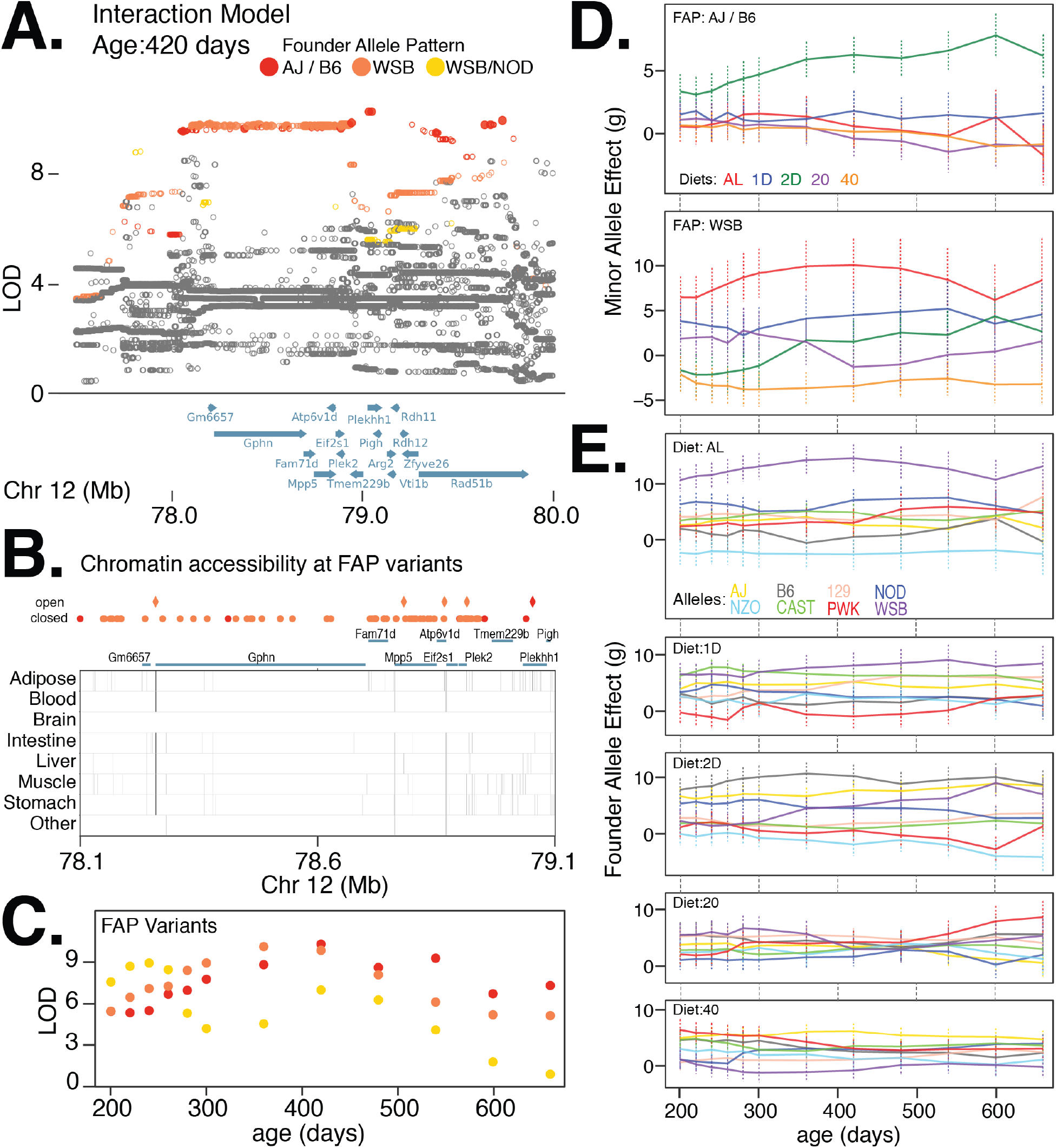
Allelic heterogeneity at a diet-dependent body weight QTL on chromosome 12. (A) Fine-mapping loci, under the interaction model, at 420 days of age. Significant variants are marked as solid circles. Colors denote variants with shared FAPs; rank 1, 2, and 3 by LOD score are colored red, orange, and yellow, respectively. (B) Significant variants, colored by their FAP, along with the gene models (shown in green) and the tissue-specific activity of regulatory elements near these variants (shown in grey). Significant variants that lie within regulatory elements are highlighted as diamonds, and regulatory elements that contain a significant variant are highlighted in dark grey. (C) Log odds ratio of association, under the interaction model, as a function of age for the lead variant from each FAP group. FAP colors are consistent with (A). (D) Diet-dependent effects (se) of the minor allele for the lead imputed variant from the AJ/B6 FAP group (top panel) and the WSB FAP group (bottom panel). (E) Diet-dependent effects (se) of the founder allele for the genotyped variant with the highest LOD score.

Despite having alleles with distinct founders-of-origin, we found that both lead FAPs had largely concordant strengths of association through age, with the WSB-specific FAP having stronger evidence at earlier ages (300-420 days) and the AJ/B6 FAP having stronger evidence at later ages (360-540 days) (Figure 6C). The minor allele of the lead imputed variant in the AJ/B6 FAP group was associated with increased body weight in the 2D fasting diet, but had little effect in the other four diets (Figure 6D). In contrast, the minor allele of the lead imputed variant from the WSB-specific FAP group had the largest positive effect on body weight in the AL and 1D fast diet and largest negative effect on body weight in the 40% CR diet (Figure 6D). Estimates of the diet-dependent effects of the AJ, B6, and WSB founder alleles at marker variants with the strongest association were consistent with the effects of FAPs predicted above: the AJ and B6 founder alleles were associated with the largest body weights in the 2D fast treatment and had little effect in the other four treatments, whereas the WSB founder allele had large, positive effects under AL and 1D fast diets and a negative effect under the 40% CR diet (Figure 6E). Taken together, these results provide evidence for allelic heterogeneity at this locus, with at least three functional alleles having distinct diet-dependent effects on body weight.

These two loci demonstrate the varied age- and diet-dependent genetic effects that shaped body weight in this DO mouse population. We found that one locus on chromosome 6 had both diet-dependent and diet-independent associations located in adjacent genomic regions and driven by different FAP groups, which is consistent with the hypothesis that the two associations are due to distinct QTLs rather than a single pleiotropic QTL. In order to determine whether this was a general feature of loci with diet-dependent and diet-independent associations, we examined eight additional such loci in our dataset (Table 1). We found six loci exhibiting a similar pattern as the chromosome 6 locus; i.e., the diet-dependent and diet-independent associations were composed of distinct FAP groups located in adjacent, distinct genomic regions (Supplemental Figure S5). The two remaining loci provided examples of a contrasting model; the diet-dependent and diet-independent associations were composed of similar (although, not identical) FAP groups centered upon the same genes (Supplemental Figure S6). The founder allele effects are distinct between the diet-dependent and diet-independent associations: the diet-dependent and diet-independent loci on chromosome 3 are due to differential effects between CAST/NOD/NZO versus B6/WSB and CAST/PWK versus B6/AJ, respectively (Supplemental Figure S6B), and the diet-dependent and diet-independent chromosome 7 loci are due to differential effects between B6/WSB/NOD versus AJ/NZO and B6/WSB/129 versus PWK (Supplemental Figure S6D). These results are consistent with the hypothesis that distinct alleles at the same locus are responsible for the diet-dependent and diet-independent associations, similar the instance of allelic heterogeneity we observed at the chromosome 12 locus (Figure 6B). Additionally, we observed one other plausible instance of allelic heterogeneity at the diet-independent QTL on chromosome 4 (Supplemental Figure S5C). In summary, fine-mapping body weight QTLs and examining their lead FAPs have revealed evidence for both allelic heterogeneity at individual loci and multiple adjacent QTLs in narrow genomic regions shaping phenotypic variation in this classic quantitative trait.

### 5.3 Nonlinear context-dependent trends in genetic effects on body weight

Given high resolution temporal measurements of body weight, we observed an age-dependent nonlinear trend in the effect of the lead variant at the diet-independent locus on chromosome 6 (Figure 5H). Conservatively, we defined age-dependent effects to be nonlinear if the trend of effect size showed a stable change in direction at any age within 30 – 660 days (e.g., an increasing effect followed by decreasing effect, each at multiple ages). Surprisingly, the nonlinear genetic effects at this locus appear to be completely driven by the AJ founder background (Figure 5J), indicating that the same allele has substantially different effects on body weight in distinct genetic backgrounds. Notably, such nonlinear trends of genetic effects with age often cannot be discerned even with large cross-sectional data that are typical of modern genome-wide association studies. To quantify the generality of such nonlinear trends, we evaluated the effect size trends at all 14 diet-independent associations identified by our study. We observed nonlinear age-dependent effects to be predominant, with 12 of the 14 loci could be classified as having nonlinear genetic effects. Additionally, nonlinearity was often specific to a subset of founder strains that are driving the associations at each locus, suggesting that the genetic background plays an important role when interpreting the dynamics of the genetic architecture of body weight in DO mice.

Along similar lines, we observed nonlinear trends with age for diet-dependent effects at the locus on chromosome 12, specifically within the WSB genetic background under the AL diet (Figure 6D). As a more general pattern, we observed such diet-specific nonlinear trends to be less common, with only 6 of 19 diet-dependent loci exhibiting nonlinearity in trends of genetic effects. In contrast to diet-independent loci, where the directionality of genetic effects were always stable across age, we found that 7 out of the 19 diet-dependent loci exhibited a switch in the directionality of effects under specific diets. One example of such a shift in genetic effect was observed in the effect of WSB-private FAP group under the 2D fasting diet (Figure 6D), where the minor allele of the lead variant in this FAP group was associated with decrease in body weight soon after the fasting intervention began, but was associated with increase in body weight about 180 days (6 months) after dietary intervention (see Supplemental Figures S8A-D for more examples).

The diet-dependent locus on chromosome 6 illustrated a rather counter-intuitive result; while we observed an approximately linear reduction in median body weight between the AL, 20% CR, and 40% CR diets in response to a linear reduction in calories (Figure 1B), the effects of this locus on body weight were nonlinear in the context of each diet (Figure 5I). We defined nonlinear diet-dependent effects as instances in which the genetic effects in the context of either the 20% CR or 1D fast treatments were substantially greater (or less) than the genetic effects in the context of AL and 40% CR diets or AL and 2D fast diets, respectively. We observed a second instance of nonlinear diet-dependent effects at a locus we mapped to chromosome 5. This locus had the strongest diet-dependent association observed in the genome. The lead FAP group, containing variants with an allele private to the NOD strain, was associated with large positive effect on body weight in the 20% CR diet, a small positive effect in the 1D fast diet, and nearly neutral effects in the 40% CR, 2D fast, and AL diets (Supplemental Figure S7). In total, we found that this pattern to be quite common, with 9 of the 19 significant loci identified under the interaction model exhibiting nonlinear diet-dependent effects (Supplemental Figure S8).

### 5.4 Fine-mapped genes implicate neurological and metabolic processes

Of the 33 significant loci from the additive and interaction models, we identified 14 loci with lead FAP groups spanning 1-3 genes (Table 1). Many of these genes implicated in modulating body weight also affect neurological behavior, consistent with the enrichment of genetic associations with body-mass index and obesity in pathways active in the central nervous system in humans [30, 31, 32,].

One example is the neuronal growth regulator 1 (Negr1) gene, a candidate linked to body weight in mid-adulthood via the lead 129/CAST/WSB FAP group within a diet-dependent locus on chromosome 3. This gene is highly expressed in the cerebral cortex and hippocampus in the rat brain [33,] and is known to regulate synapse formation of hippocampal neurons and promote neurite outgrowth of cortical neurons [34, 35,]. Negr1 has also been implicated in obesity [36,], autistic behavior, memory deficits, and increased susceptibility to seizures [37,] in mice, and body-mass index [38,] and major depressive disorder [39,] in humans. Another example is the gephyrin gene (Gphn) implicated by two distinct FAP groups in the diet-dependent locus on chromosome 12 (Figure 6A). Gephyrin is a key structural protein at neuronal synapses that ensures proper localization of postsynaptic inhibitory receptors. Gephyrin is also known to physically interact with mTOR and is required for mTOR signaling [40,], suggesting two plausible pathways for influencing body weight. On chromosome 1, murine Trmt1l (Trm1-like), a gene with sequence similarity to orthologous tRNA methyltransferases in other species, was linked to body weight during early development (Supplemental Figure S5A). Mice deficient in this gene, while viable, have been found to exhibit altered motor coordination and abnormal exploratory behavior [41,], suggesting that the association at this locus is possibly mediated through modulating exploratory behavior.

Among candidates that affect metabolic processes, Creb5, a gene linked to diet-independent effects on body weight (Figure 5B), has previously been reported to be linked to metabolic phenotypes in humans, with differential DNA methylation detected between individuals with large differences in waist circumference, hypercholesterolemia, and metabolic syndrome [42,]. Another gene important for metabolic control in the liver, PI3K-C2*γ* was linked to diet-dependent effects on body weight in early adulthood. PI3K-C2*γ*-deficient mice are known to exhibit reduced liver accumulation of glycogen and develop hyperlipidemia, adiposity, and insulin resistance with age [43,]. Edem3, another candidate gene at the locus on chromosome 1 linked to body weight in early development (Supplemental Figure S5A), has previously been linked to short stature in humans based on family-based exome sequencing and differential expression in chondrocytes [44,]. Possibly sharing a similar mechanism, Adam12, a candidate gene in both a diet-independent and diet-dependent locus on chromosome 7, is known to play an important role in the differentiation, proliferation, and maturation of chondrocytes [45,].

Thus, our fine-mapping strategy using FAP groups has highlighted several candidate genes associated with body weight, many of which are known to play important roles in a range of processes including metabolism, skeletal growth, motor coordination, and behavior.

## 6 Future Considerations

To summarize, we found that the effects of age and diet on body weight differ substantially with respect to the genetic background and type of dietary intervention imposed. These results highlight that with knowledge of these environment-dependent effects, we can generate more accurate predictions of body weight trajectories than would be possible from knowledge of genotype, age, or diet alone. Moreover, with the elucidation of specific candidate genes and variants underlying these effects, we are not limited to predicting how this complex quantitative trait changes with age, but can also identify specific targets for genetic or pharmacological manipulation in an effort to improve organismal health.

One important limitation to our study is the lack of direct measurements of food consumption and feeding behaviors for each mouse in the population; this makes it difficult to ascertain how much variation in body weight can be ascribed to variation in such behaviors. Additionally, caloric restriction was imposed based on the average food consumed by a typical DO mouse rather than a per-mouse baseline of food consumption. Furthermore, caloric restriction interventions were imposed on a per-cage basis, not a per-mouse basis, because all animals were housed in groups of eight. Therefore, the social hierarchy within each cage likely contributed to additional variation in body weight [46,]. Accounting for these sources of variation will be a promising avenue for future research, helping interpret many of the associations identified in our study.

A second important consideration for this study is the potential for survival bias to lead to inflated PVE values and false positive associations at later ages. To evaluate the presence of survival bias, we computed PVE at all ages restricting to animals that survived to 660 days of age (75% of animals in our study). We observed similar PVE values to those from the full data set, across the full age range, suggesting that survival bias has very little effect on the results presented in this paper (Supplemental Figure S3C, top left panel). However, as these animals age and the survival bias of the population increases, genetic analyses of body weight at ages past 660 days will need to explicitly account for this effect.

In this study, we have elucidated the dynamics and context-dependence of the genetic architecture of body weight from 60 to 660 days of age. By 660 days of age, nearly all surviving animals have realized their maximum adult body weight and the majority of animals have yet to experience appreciable loss of body weight indicative of late-age physiological decline. Under AL diet, reduced body weight has been associated with greater longevity, whereas under 40% CR conditions, greater longevity is associated with the maintenance of high body weight [47,]. Our future research will assess whether we observe a similar result in this DO population and determine whether alleles at body weight loci are predictive of lifespan in a diet-dependent manner.

## 7 Appendix A: Proportion of variance explained

Decomposing the phenotype into genetic and non-genetic effects, **Y** = **Y**_**G**_ +**Y**_*ε*_, the expected proportion of phenotypic variance explained by genetic effects is approximately given as

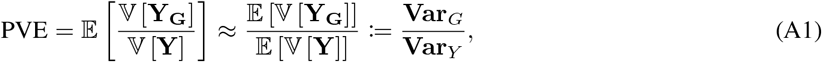

where 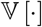 denotes the sample variance. The expected sample phenotypic variance conditional on environment *e* can be written as

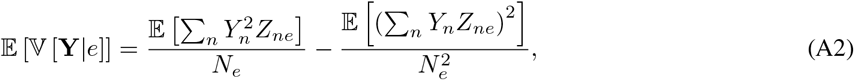

where *Z*_*ne*_ is an indicator variable denoting whether sample *n* belongs to environment *e* and *N*_*e*_ = ∑_*n*_ *Z*_*ne*_ is the number of samples in environment *e*. Under the GxEMM model, starting from Equation 1 and integrating out the random effects, we can write the numerator of the first term in the expectation as

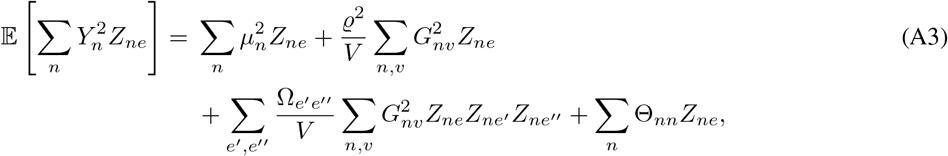

and the numerator of the second term in the expectation as

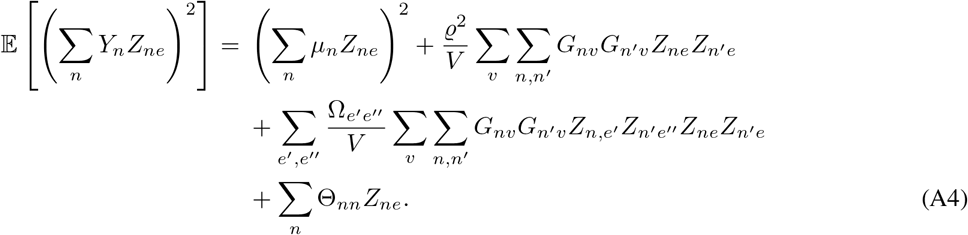

Therefore, the expected sample phenotypic variance can be decomposed as follows

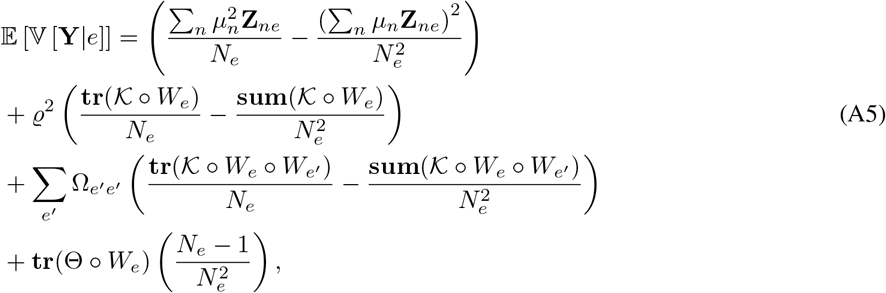

where **tr**(·) denotes that trace of a matrix, **sum**(·) denotes the sum of all entries of the matrix, and *A* ◦ *B* denotes the Hadamard product of matrices *A* and *B*. The first term quantifies the phenotypic variance explained by fixed effects, the second and third terms together quantify the phenotypic variance explained by genetic effects 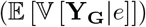, and the fourth term quantifies the residual (unexplained) phenotypic variance. The proportion of variance explained by genetics conditional on environment can now be computed using Equation A1.

The expected total sample phenotypic variance (across all environments) again has two terms, as in Equation A2; the numerator of the first term is written as

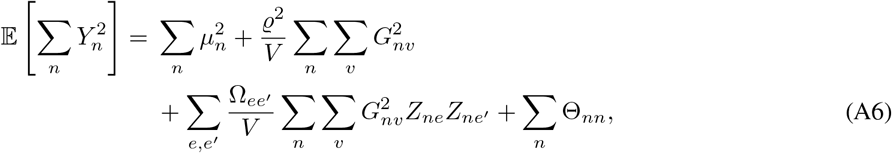

and the numerator of the second term is written as

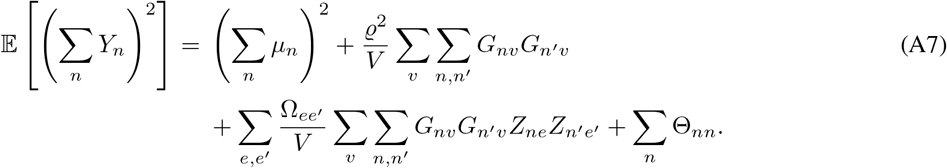

The expected total sample phenotypic variance can be decomposed as follows

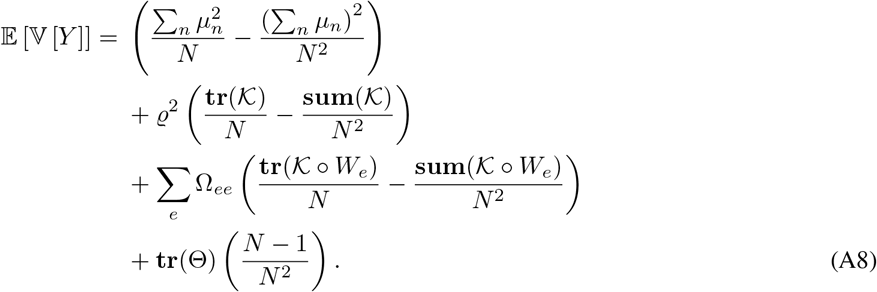

The total proportion of variance explained by genetics in the entire sample can be computed by substituting the above in Equation A1.

Under the EMMA model, the expected total sample phenotypic variance simplifies to

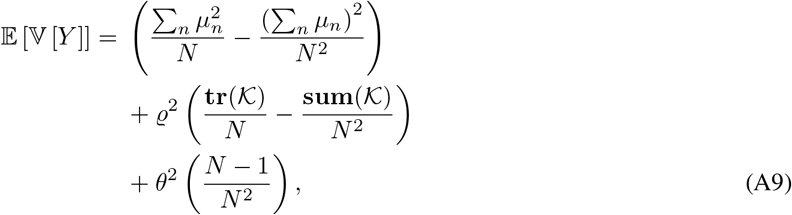

where *θ* denotes the homoscedastic noise. The first component quantifies the phenotypic variance explained by fixed effects, the second component quantifies the phenotypic variance explained by genetic effects 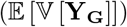, and the third component quantifies the residual (unexplained) phenotypic variance. Substituting these into Equation A1 gives us the proportion of variance explained by genetics under the EMMA model.

## 8 Appendix B: Likelihood and gradients for GxEMM model

Under the full GxEMM model, the phenotype **Y** depends on fixed and random effects as follows:

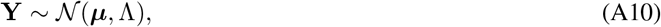

where ***μ*** = *α*_0_ + ∑_*e*_ **Z**_*e*_*α*_*e*_ captures all fixed effects and 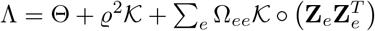 captures the covariance after integrating out the random effects. The parameters to be estimated in this model are ***α***, ***σ***, ϱ, and Ω. The complete log likelihood can be written as

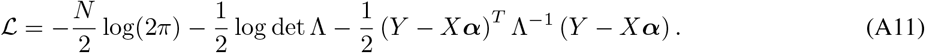

Maximizing the log likelihood over ***α*** gives us

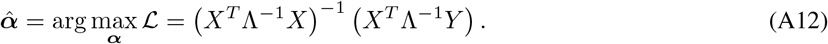

Substituting this into the log likelihood, we get

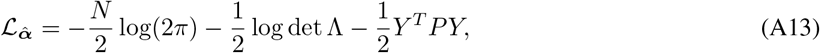

where *P* = Λ^−1^ − Λ^−1^*X*(*X*^*T*^ Λ^−1^*X*)^−1^ *X*^*T*^ Λ^−1^.

Computing the gradient of 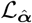 involves evaluating the following gradients,

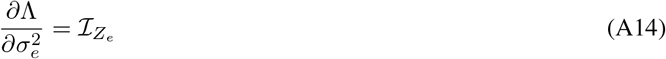

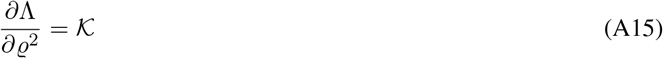

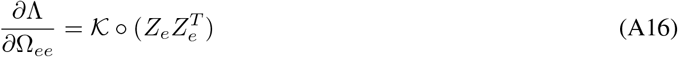

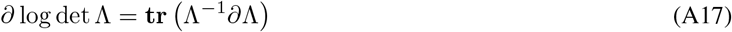

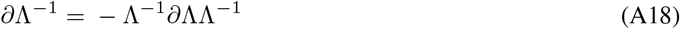

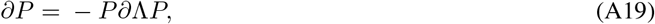

where 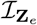 is a diagonal matrix with elements of the vector **Z**_*e*_ on the diagonal. Thus,

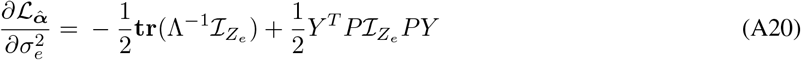

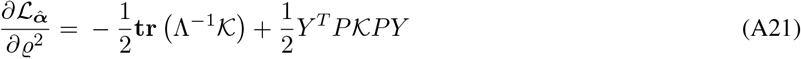

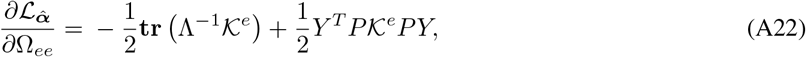

where 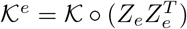

## 9 Supplemental Figures

**Figure S1:**
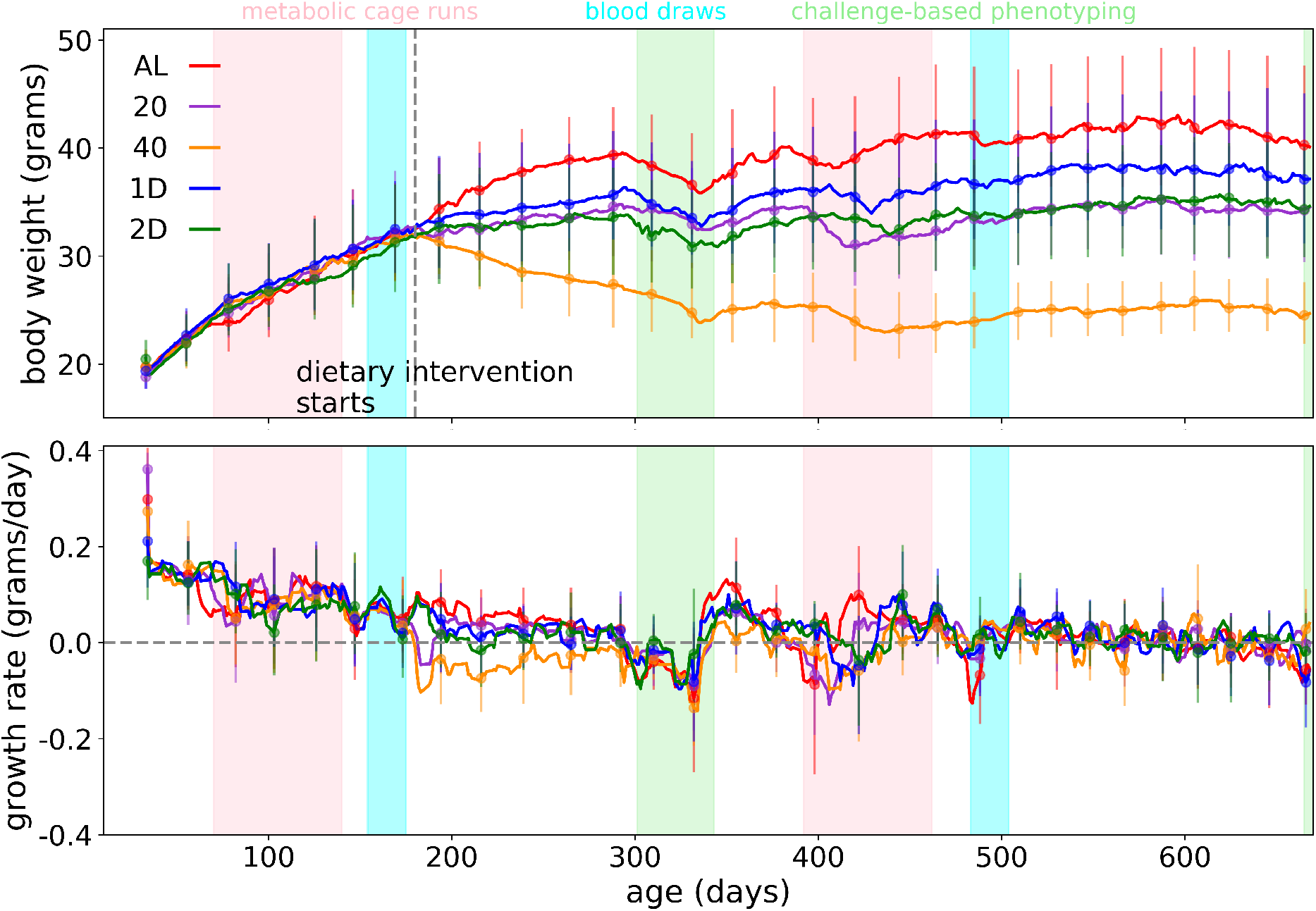
Raw measurements. (A) Body weight and (B) growth rate trends as a function of age. Mice in this study also undering an array of phenotyping procedures; the age range for metabolic cage phenotyping is highlighted in pink, the age range for blood draws is highlighted in blue, and the age range for a battery of challenge-based phenotyping procedures is highlighted in green.

**Figure S2:**
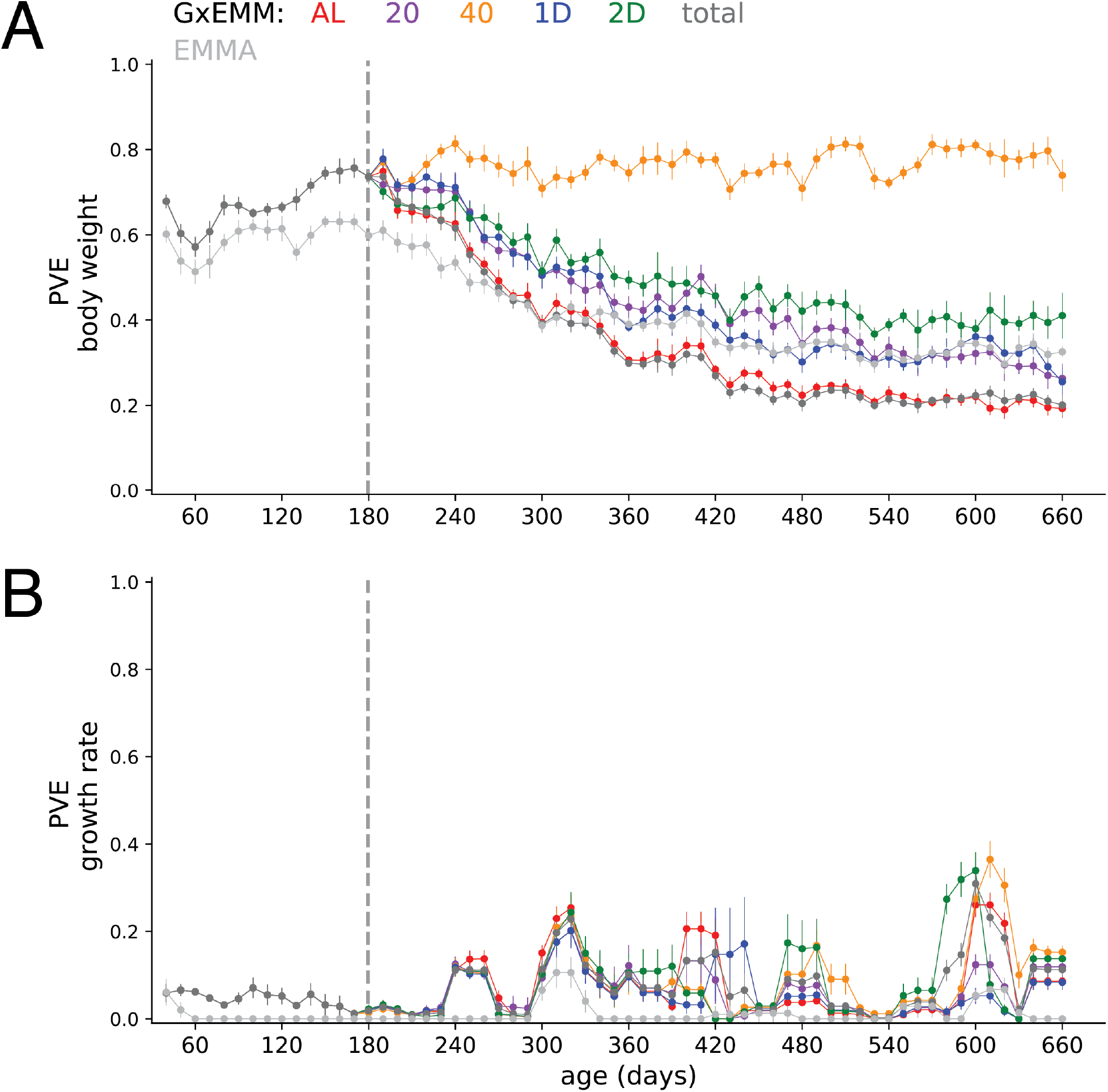
PVE using raw measurements. PVE of (A) Body weight and (B) growth rate, estimated using raw body weight measurements.

**Figure S3:**
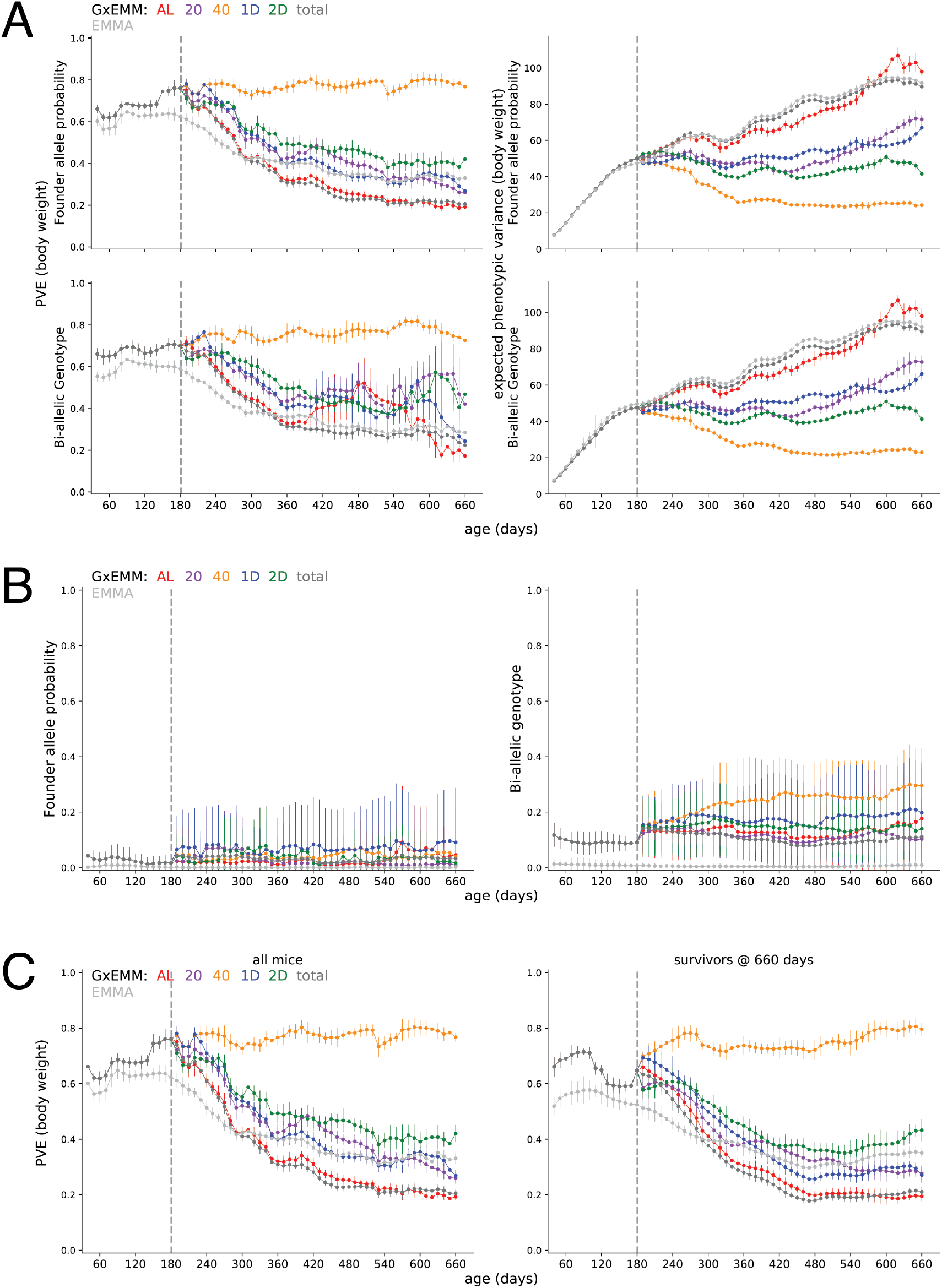
Phenotypic variance explained by genetics. (A) PVE (left column) and expected phenotypic variance (right column) estimated using kinship calculated from founder-of-origin allele probabilities (top row) and bi-allelic genotypes (bottom row) (B) PVE estimated using kinship calculated from founder-of-origin allele probabilities (left column) and bi-allelic genotypes (right column), after randomly permuting body weight trajectories across mice, within each dietary intervention. (C) PVE estimated using all mice in the study (left column) and mice that survived to 660 days of age (right column).

**Figure S4:**
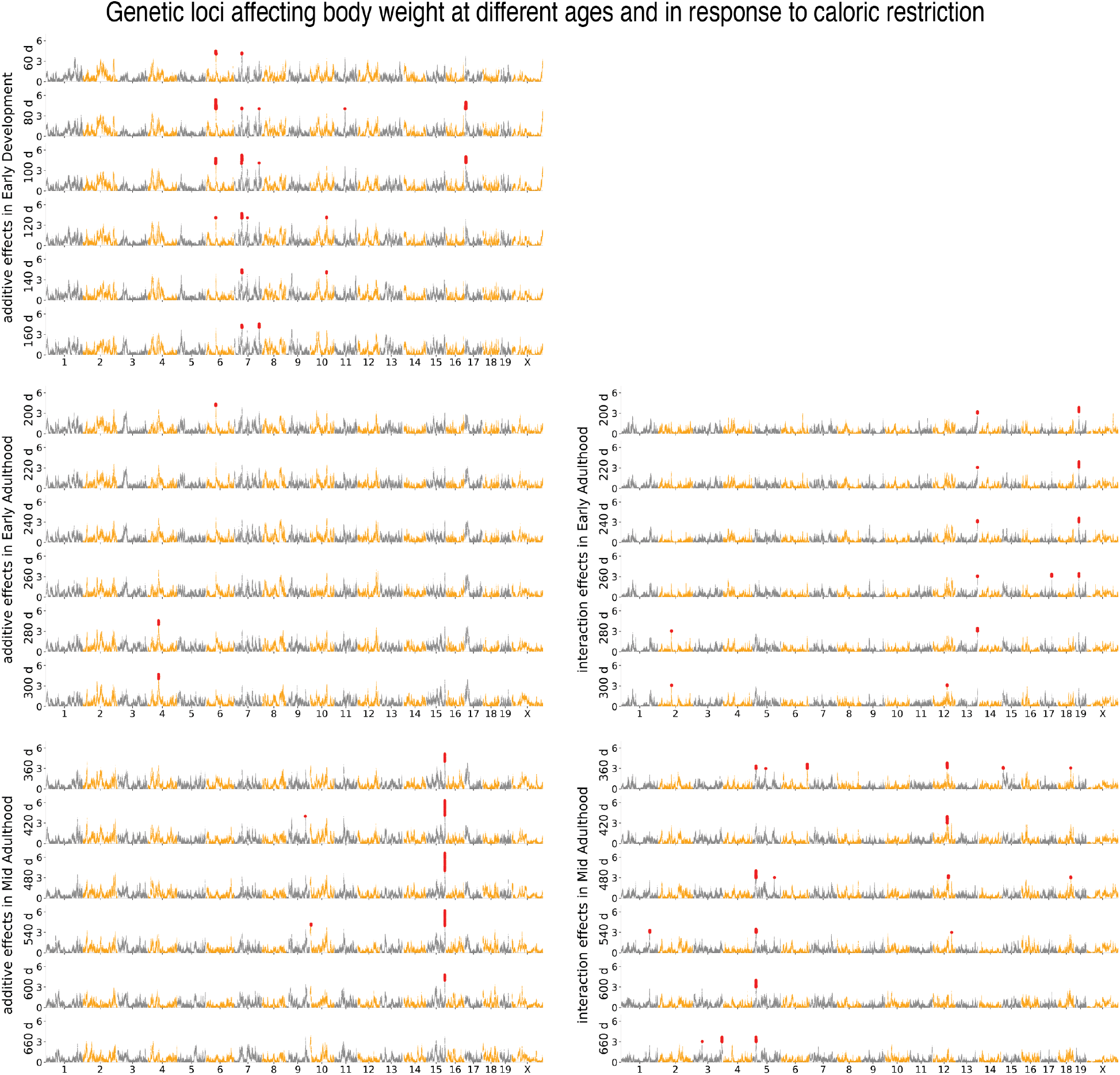
Age- and diet-dependent Manhattan plots for body weight. Genetic loci associated with body weight at different ages identified under the additive genetic model (subpanels on the left) and genotype-diet interaction model (subpanels on the right). Each circle is a genotyped marker, significant markers are in red.

**Figure S5:**
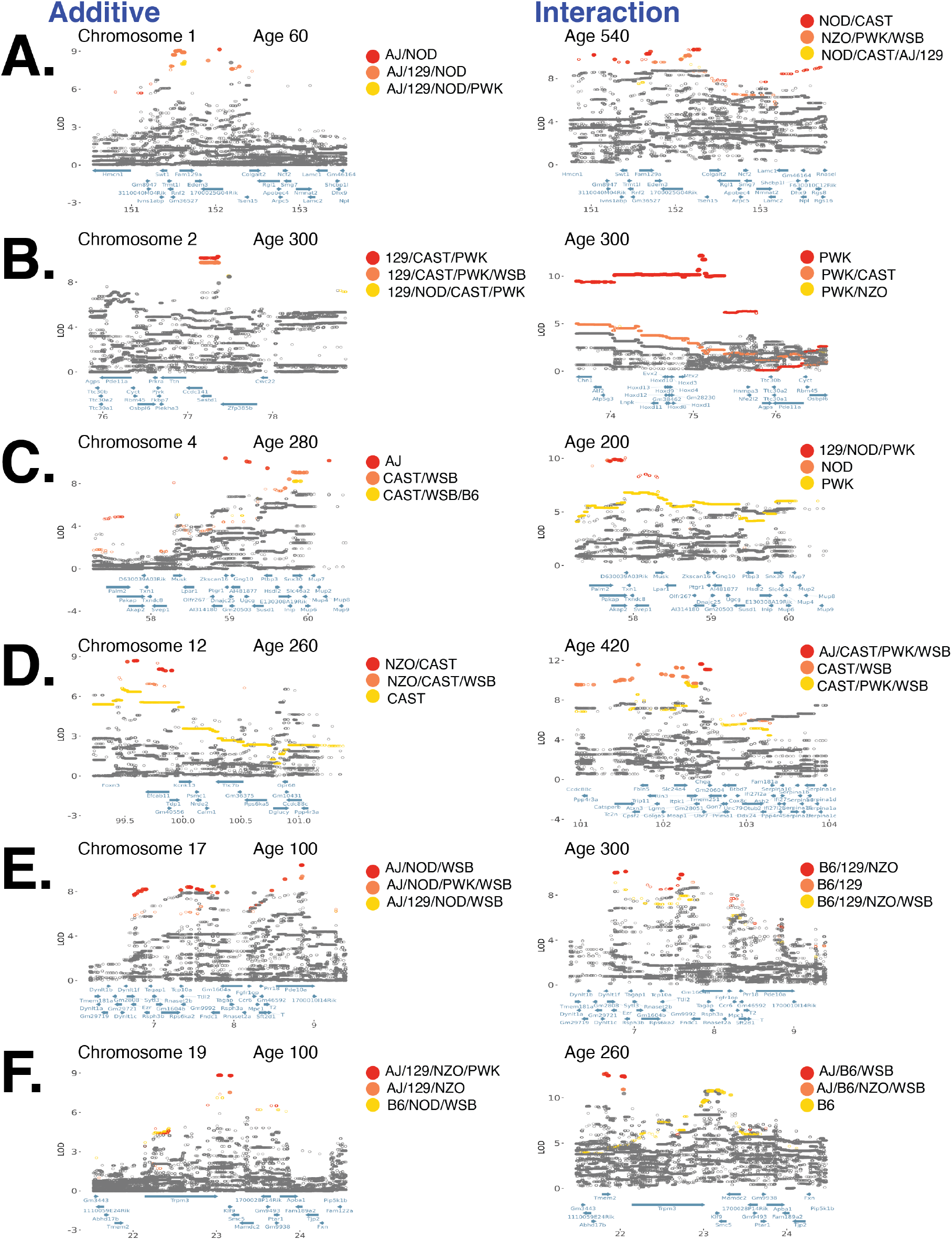
Loci with significant diet-independent and diet-dependent associations with body weight. (A) Fine-mapping a locus on chromosome 1 associated with body weight under the additive model (left column) and genotype-diet interaction model (right column). Each circle is a bi-allelic variant, both imputed and genotyped, and solid circles denote significantly associated variants. Variants are colored according to their FAP; FAPs of rank 1, 2, and 3 (based on LOD score) are colored red, orange, and yellow, respectively. Panels (B) - (F) are the same as (A) for loci on chromosomes: 2, 4, 12, 17, and 19.

**Figure S6:**
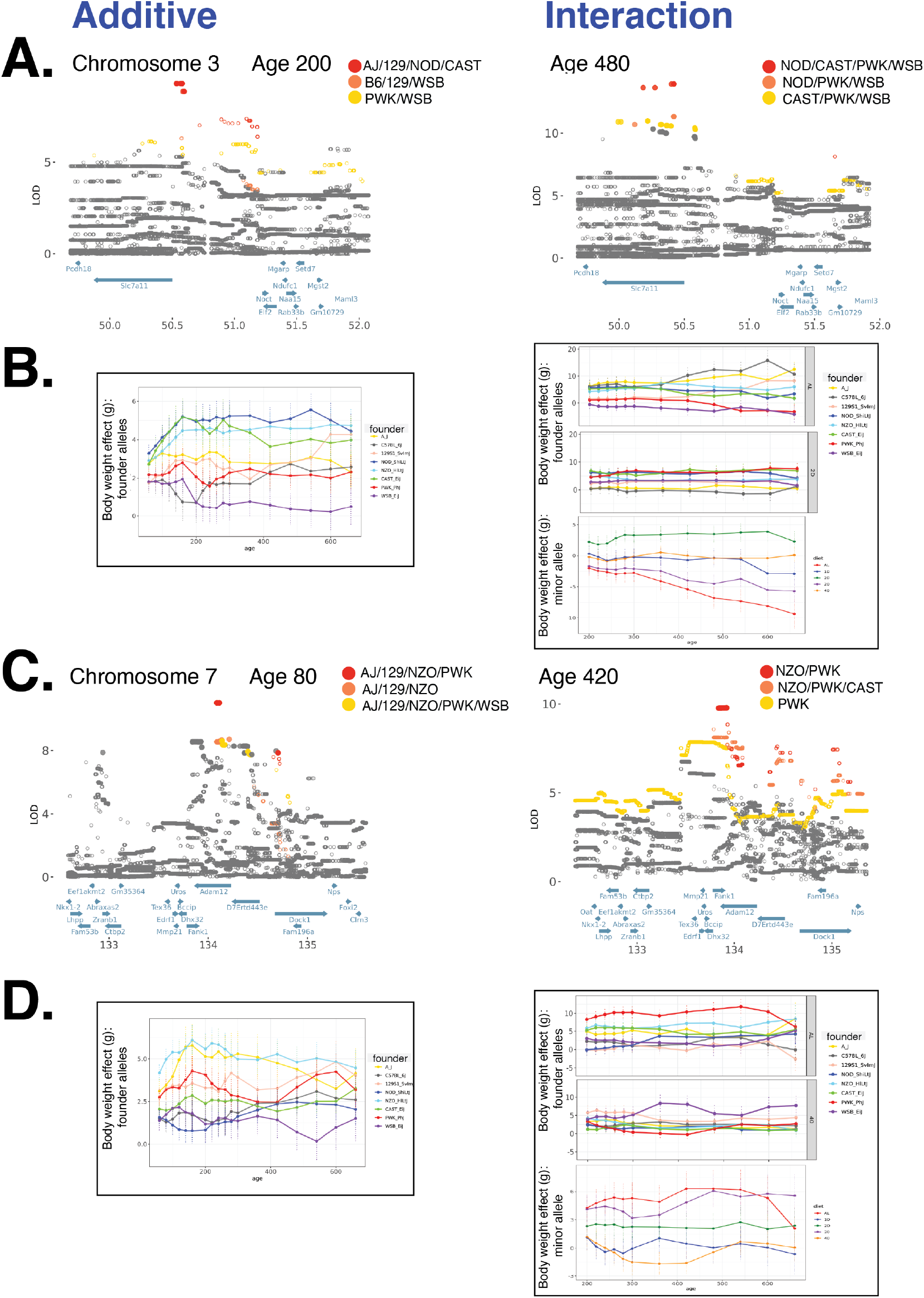
Allelic heterogeneity at loci with significant diet-independent and diet-dependent associations with body weight. (A) Fine-mapping a locus on chromosome 3 associated with body weight under the additive model (left column) and genotype-diet interaction model (right column). Each circle is a bi-allelic variant, both imputed and genotyped, and solid circles denote significantly associated variants. Variants are colored according to their FAP; FAP with ranks 1, 2, and 3 are colored red, orange, and yellow, respectively. (B) Founder allele effect as a function of age for the lead FAP variant from (A) for the diet-independent association (left column) and the diet-dependent association (right column). Panels (C) and (D) are the same as (A) and (B) for a locus on chromosome 7.

**Figure S7:**
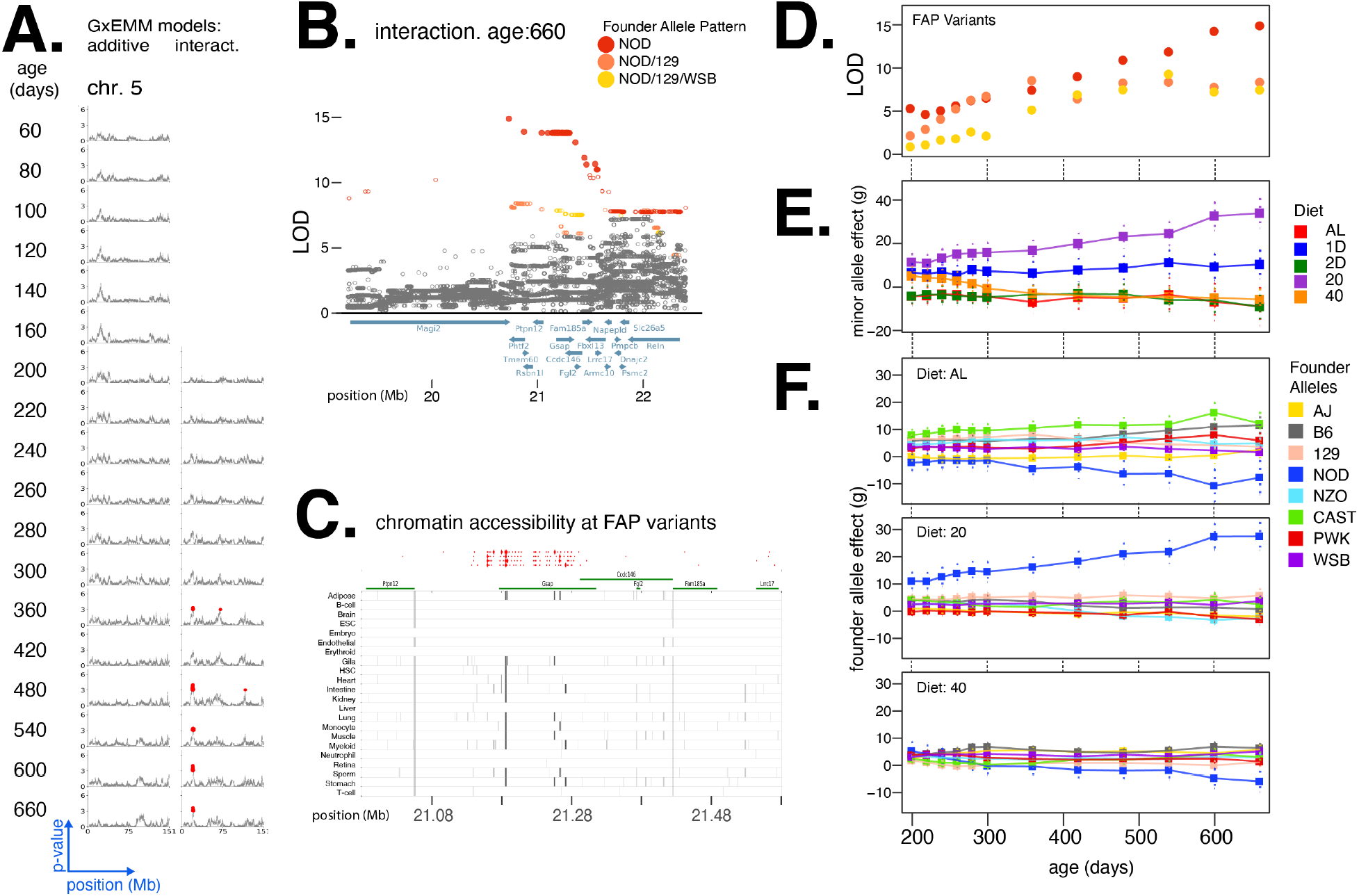
Diet-dependent association with body weight in a locus on chromosome 5. (A) Manhattan plots of additive genetic associations and genotype-diet associations on chromosome 5 at multiple ages. (B) Fine-mapping a locus associated with body weight in diet-dependent manner at 660 days of age. Each circle is a bi-allelic variant, both imputed and genotyped and solid circles denote significantly associated variants. Variants are colored according to their FAP; FAPs of rank 1, 2, and 3 (by LOD score) are colored red, orange, and yellow, respectively. (C) Significant variants, colored by their FAP, along with the gene models (shown in green) and the tissue-specific activity of regulatory elements near these variants (shown in grey). Significant variants that lie within regulatory elements are highlighted as diamonds, and regulatory elements that contain a significant variant are highlighted in dark grey. (D) Log odds ratio as a function of age, for a single variant that exhibits the strongest association from each of the top three FAPs. (E) Diet-dependent effect size of the minor allele as a function of age, for the variant with the strongest association. (F) Estimated effect of each founder allele in three diets (AL, 20, 40), at the genotyped variant with the strongest diet-dependent association.

**Figure S8:**
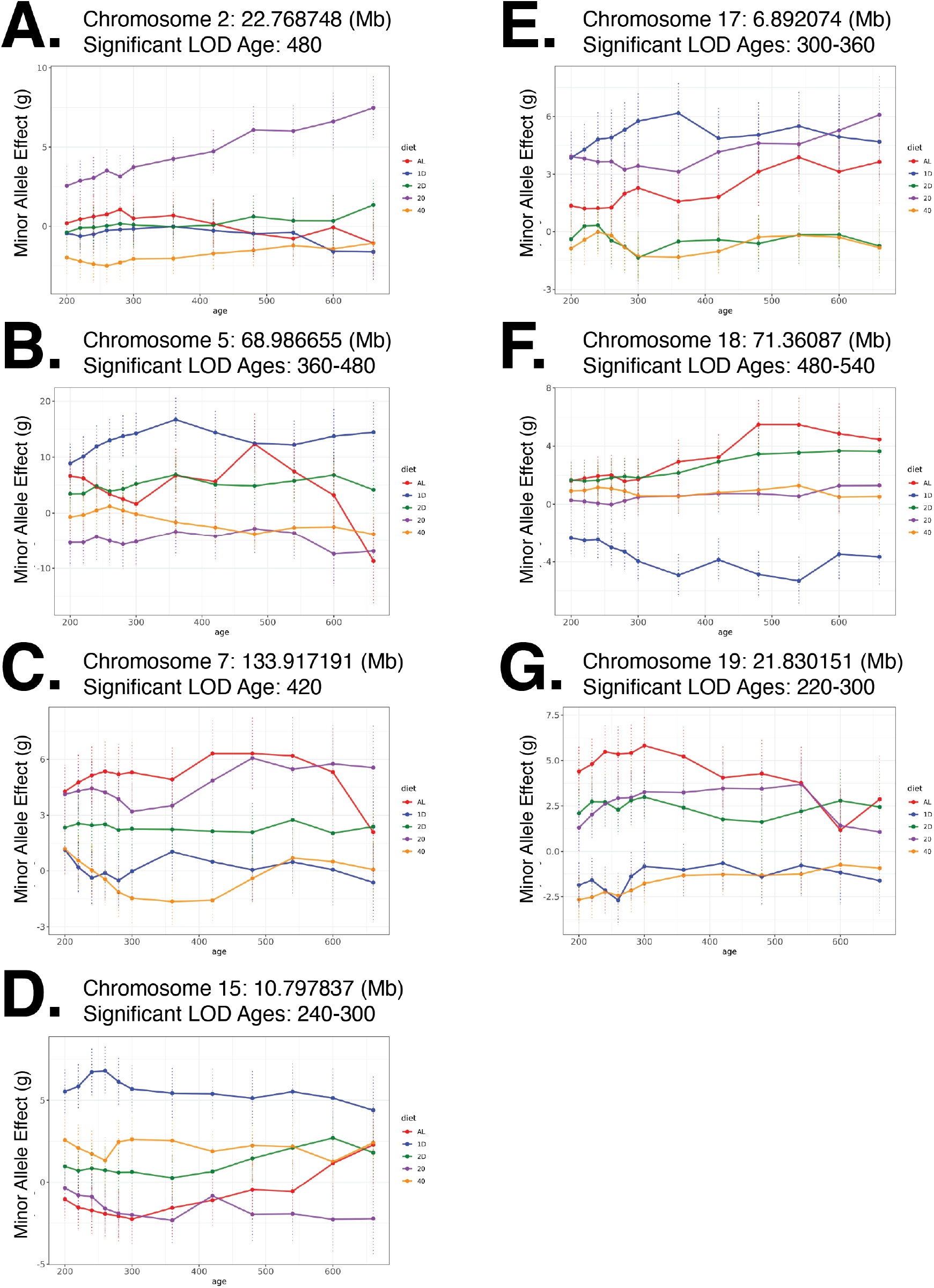
Nonlinear trends in genetic effects with respect to age and dietary intervention. For each fine-mapped locus, we note the location of the variant with the strongest association and the ages at which the genetic association is significant, and plot the estimated effect (SE) of this variant as a function of age, under different diets.

## References

[1] Matthew R Robinson, Geoffrey English, Gerhard Moser, Luke R Lloyd-Jones, Marcus A Triplett, Zhihong Zhu, Ilja M Nolte, Jana V van Vliet-Ostaptchouk, Harold Snieder, LifeLines Cohort Study, Tonu Esko, Lili Milani, Reedik Mägi, Andres Metspalu, Patrik K E Magnusson, Nancy L Pedersen, Erik Ingelsson, Magnus Johannesson, Jian Yang, David Cesarini, and Peter M Visscher. Genotype-covariate interaction effects and the heritability of adult body mass index. Nat. Genet., 49(8):1174–1181, August 2017.

[2] Jae Hoon Sul, Michael Bilow, Wen-Yun Yang, Emrah Kostem, Nick Furlotte, Dan He, and Eleazar Eskin. Accounting for population structure in Gene-by-Environment interactions in Genome-Wide association studies using mixed models. PLoS Genet., 12(3):e1005849, March 2016.

[3] Rachel Moore, Francesco Paolo Casale, Marc Jan Bonder, Danilo Horta, BIOS Consortium, Lude Franke, Inês Barroso, and Oliver Stegle. A linear mixed-model approach to study multivariate gene-environment interactions. Nat. Genet., 51(1):180–186, January 2019.

[4] Daniel E Runcie and Lorin Crawford. Fast and flexible linear mixed models for genome-wide genetics. PLoS Genet., 15(2):e1007978, February 2019.

[5] Andy Dahl, Khiem Nguyen, Na Cai, Michael J Gandal, Jonathan Flint, and Noah Zaitlen. A robust method uncovers significant Context-Specific heritability in diverse complex traits. Am. J. Hum. Genet., 106(1):71–91, January 2020.

[6] J M Cheverud, E J Routman, F A Duarte, B van Swinderen, K Cothran, and C Perel. Quantitative trait loci for murine growth. Genetics, 142(4):1305–1319, April 1996.

[7] B Riska, W R Atchley, and J J Rutledge. A genetic analysis of targeted growth in mice. Genetics, 107(1):79–101, May 1984.

[8] Melissa M Gray, Michelle D Parmenter, Caley A Hogan, Irene Ford, Richard J Cuthbert, Peter G Ryan, Karl W Broman, and Bret A Payseur. Genetics of rapid and extreme size evolution in island mice. Genetics, 201(1):213–228, September 2015.

[9] Clarissa C Parker, Shyam Gopalakrishnan, Peter Carbonetto, Natalia M Gonzales, Emily Leung, Yeonhee J Park, Emmanuel Aryee, Joe Davis, David A Blizard, Cheryl L Ackert-Bicknell, Arimantas Lionikas, Jonathan K Pritchard, and Abraham A Palmer. Genome-wide association study of behavioral, physiological and gene expression traits in outbred CFW mice. Nat. Genet., 48(8):919–926, August 2016.

[10] Brad A Rikke, Matthew E Battaglia, David B Allison, and Thomas E Johnson. Murine weight loss exhibits significant genetic variation during dietary restriction. Physiol. Genomics, 27(2):122–130, October 2006.

[11] Artem Vorobyev, Yask Gupta, Tanya Sezin, Hiroshi Koga, Yannic C Bartsch, Meriem Belheouane, Sven Künzel, Christian Sina, Paul Schilf, Heiko Körber-Ahrens, Foteini Beltsiou, Anna Lara Ernst, Stanislav Khil’chenko, Hassanin Al-Aasam, Rudolf A Manz, Sandra Diehl, Moritz Steinhaus, Joanna Jascholt, Phillip Kouki, Wolf-Henning Boehncke, Tanya N Mayadas, Detlef Zillikens, Christian D Sadik, Hiroshi Nishi, Marc Ehlers, Steffen Möller, Katja Bieber, John F Baines, Saleh M Ibrahim, and Ralf J Ludwig. Gene-diet interactions associated with complex trait variation in an advanced intercross outbred mouse line. Nat. Commun., 10(1):4097, September 2019.

[12] Alexessander Couto Alves, N Maneka G De Silva, Ville Karhunen, Ulla Sovio, Shikta Das, H Rob Taal, Nicole M Warrington, Alexandra M Lewin, Marika Kaakinen, Diana L Cousminer, Elisabeth Thiering, Nicholas J Timpson, Tom A Bond, Estelle Lowry, Christopher D Brown, Xavier Estivill, Virpi Lindi, Jonathan P Bradfield, Frank Geller, Doug Speed, Lachlan J M Coin, Marie Loh, Sheila J Barton, Lawrence J Beilin, Hans Bisgaard, Klaus Bønnelykke, Rohia Alili, Ida J Hatoum, Katharina Schramm, Rufus Cartwright, Marie-Aline Charles, Vincenzo Salerno, Karine Clément, Annique A J Claringbould, BIOS Consortium, Cornelia M van Duijn, Elena Moltchanova, Johan G Eriksson, Cathy Elks, Bjarke Feenstra, Claudia Flexeder, Stephen Franks, Timothy M Frayling, Rachel M Freathy, Paul Elliott, Elisabeth Widén, Hakon Hakonarson, Andrew T Hattersley, Alina Rodriguez, Marco Banterle, Joachim Heinrich, Barbara Heude, John W Holloway, Albert Hofman, Elina Hyppönen, Hazel Inskip, Lee M Kaplan, Asa K Hedman, Esa Läärä, Holger Prokisch, Harald Grallert, Timo A Lakka, Debbie A Lawlor, Mads Melbye, Tarunveer S Ahluwalia, Marcella Marinelli, Iona Y Millwood, Lyle J Palmer, Craig E Pennell, John R Perry, Susan M Ring, Markku J Savolainen, Fernando Rivadeneira, Marie Standl, Jordi Sunyer, Carla M T Tiesler, Andre G Uitterlinden, William Schierding, Justin M O’Sullivan, Inga Prokopenko, Karl-Heinz Herzig, George Davey Smith, Paul O’Reilly, Janine F Felix, Jessica L Buxton, Alexandra I F Blakemore, Ken K Ong, Vincent W V Jaddoe, Struan F A Grant, Sylvain Sebert, Mark I McCarthy, Marjo-Riitta Järvelin, and Early Growth Genetics (EGG) Consortium. GWAS on longitudinal growth traits reveals different genetic factors influencing infant, child, and adult BMI. Sci Adv, 5(9):eaaw3095, September 2019.

[13] Huanwei Wang, Futao Zhang, Jian Zeng, Yang Wu, Kathryn E Kemper, Angli Xue, Min Zhang, Joseph E Powell, Michael E Goddard, Naomi R Wray, Peter M Visscher, Allan F McRae, and Jian Yang. Genotype-by-environment interactions inferred from genetic effects on phenotypic variability in the UK biobank. Sci Adv, 5(8):eaaw3538, August 2019.

[14] Karen L Svenson, Daniel M Gatti, William Valdar, Catherine E Welsh, Riyan Cheng, Elissa J Chesler, Abraham A Palmer, Leonard McMillan, and Gary A Churchill. High-resolution genetic mapping using the mouse diversity outbred population. Genetics, 190(2):437–447, February 2012.

[15] Seung-Jean Kim, Kwangmoo Koh, Stephen Boyd, and Dimitry Gorinevsky. *ℓ*_1_ trend filtering. SIAM Rev., 51(2):339–360, May 2009.

[16] Andrew P Morgan, Chen-Ping Fu, Chia-Yu Kao, Catherine E Welsh, John P Didion, Liran Yadgary, Leeanna Hyacinth, Martin T Ferris, Timothy A Bell, Darla R Miller, Paola Giusti-Rodriguez, Randal J Nonneman, Kevin D Cook, Jason K Whitmire, Lisa E Gralinski, Mark Keller, Alan D Attie, Gary A Churchill, Petko Petkov, Patrick F Sullivan, Jennifer R Brennan, Leonard McMillan, and Fernando Pardo-Manuel de Villena. The mouse universal genotyping array: From substrains to subspecies. G3, 6(2):263–279, December 2015.

[17] Karl W Broman, Daniel M Gatti, Petr Simecek, Nicholas A Furlotte, Pjotr Prins, Śaunak Sen, Brian S Yandell, and Gary A Churchill. R/qtl2: Software for mapping quantitative trait loci with High-Dimensional data and multiparent populations. Genetics, 211(2):495–502, February 2019.

[18] S E Lincoln and E S Lander. Systematic detection of errors in genetic linkage data. Genomics, 14(3):604–610, November 1992.

[19] Hyun Min Kang, Noah A Zaitlen, Claire M Wade, Andrew Kirby, David Heckerman, Mark J Daly, and Eleazar Eskin. Efficient control of population structure in model organism association mapping. Genetics, 178(3):1709–1723, March 2008.

[20] Christoph Lippert, Jennifer Listgarten, Ying Liu, Carl M Kadie, Robert I Davidson, and David Heckerman. FaST linear mixed models for genome-wide association studies. Nat. Methods, 8(10):833–835, September 2011.

[21] Xiang Zhou and Matthew Stephens. Genome-wide efficient mixed-model analysis for association studies. Nat. Genet., 44(7):821–824, June 2012.

[22] Julian Besag and Peter Clifford. Sequential monte carlo p-values. Biometrika, 78(2):301–304, 1991.

[23] Heejung Shim and Matthew Stephens. Wavelet-based genetic association analysis of functional phenotypes arising from high-throughput sequencing assays. Ann. Appl. Stat., 9(2):655–686, 2015.

[24] Mark Abney. Permutation testing in the presence of polygenic variation. Genet. Epidemiol., 39(4):249–258, May 2015.

[25] David L Aylor, William Valdar, Wendy Foulds-Mathes, Ryan J Buus, Ricardo A Verdugo, Ralph S Baric, Martin T Ferris, Jeff A Frelinger, Mark Heise, Matt B Frieman, Lisa E Gralinski, Timothy A Bell, John D Didion, Kunjie Hua, Derrick L Nehrenberg, Christine L Powell, Jill Steigerwalt, Yuying Xie, Samir N P Kelada, Francis S Collins, Ivana V Yang, David A Schwartz, Lisa A Branstetter, Elissa J Chesler, Darla R Miller, Jason Spence, Eric Yi Liu, Leonard McMillan, Abhishek Sarkar, Jeremy Wang, Wei Wang, Qi Zhang, Karl W Broman, Ron Korstanje, Caroline Durrant, Richard Mott, Fuad A Iraqi, Daniel Pomp, David Threadgill, Fernando Pardo-Manuel de Villena, and Gary A Churchill. Genetic analysis of complex traits in the emerging collaborative cross. Genome Res., 21(8):1213–1222, August 2011.

[26] Thomas M Keane, Leo Goodstadt, Petr Danecek, Michael A White, Kim Wong, Binnaz Yalcin, Andreas Heger, Avigail Agam, Guy Slater, Martin Goodson, Nicholas A Furlotte, Eleazar Eskin, Christoffer Nellåker, Helen Whitley, James Cleak, Deborah Janowitz, Polinka Hernandez-Pliego, Andrew Edwards, T Grant Belgard, Peter L Oliver, Rebecca E McIntyre, Amarjit Bhomra, Jérôme Nicod, Xiangchao Gan, Wei Yuan, Louise van der Weyden, Charles A Steward, Sendu Bala, Jim Stalker, Richard Mott, Richard Durbin, Ian J Jackson, Anne Czechanski, José Afonso Guerra-Assunção, Leah Rae Donahue, Laura G Reinholdt, Bret A Payseur, Chris P Ponting, Ewan Birney, Jonathan Flint, and David J Adams. Mouse genomic variation and its effect on phenotypes and gene regulation. Nature, 477(7364):289–294, September 2011.

[27] Abanish Singh. Allele heterogeneity. In Marc D Gellman and J Rick Turner, editors, Encyclopedia of Behavioral Medicine, pages 66–67. Springer New York, New York, NY, 2013.

[28] Darren A Cusanovich, Andrew J Hill, Delasa Aghamirzaie, Riza M Daza, Hannah A Pliner, Joel B Berletch, Galina N Filippova, Xingfan Huang, Lena Christiansen, William S DeWitt, Choli Lee, Samuel G Regalado, David F Read, Frank J Steemers, Christine M Disteche, Cole Trapnell, and Jay Shendure. A Single-Cell atlas of in vivo mammalian chromatin accessibility. Cell, 174(5):1309–1324.e18, August 2018.

[29] David U Gorkin, Iros Barozzi, Yuan Zhao, Yanxiao Zhang, Hui Huang, Ah Young Lee, Bin Li, Joshua Chiou, Andre Wildberg, Bo Ding, Bo Zhang, Mengchi Wang, J Seth Strattan, Jean M Davidson, Yunjiang Qiu, Veena Afzal, Jennifer A Akiyama, Ingrid Plajzer-Frick, Catherine S Novak, Momoe Kato, Tyler H Garvin, Quan T Pham, Anne N Harrington, Brandon J Mannion, Elizabeth A Lee, Yoko Fukuda-Yuzawa, Yupeng He, Sebastian Preissl, Sora Chee, Jee Yun Han, Brian A Williams, Diane Trout, Henry Amrhein, Hongbo Yang, J Michael Cherry, Wei Wang, Kyle Gaulton, Joseph R Ecker, Yin Shen, Diane E Dickel, Axel Visel, Len A Pennacchio, and Bing Ren. An atlas of dynamic chromatin landscapes in mouse fetal development. Nature, 583(7818):744–751, July 2020.

[30] Adam E Locke, Bratati Kahali, Sonja I Berndt, Anne E Justice, Tune H Pers, Felix R Day, Corey Powell, Sailaja Vedantam, Martin L Buchkovich, Jian Yang, Damien C Croteau-Chonka, Tonu Esko, Tove Fall, Teresa Ferreira, Stefan Gustafsson, Zoltán Kutalik, Jian’an Luan, Reedik Mägi, Joshua C Randall, Thomas W Winkler, Andrew R Wood, Tsegaselassie Workalemahu, Jessica D Faul, Jennifer A Smith, Jing Hua Zhao, Wei Zhao, Jin Chen, Rudolf Fehrmann, Åsa K Hedman, Juha Karjalainen, Ellen M Schmidt, Devin Absher, Najaf Amin, Denise Anderson, Marian Beekman, Jennifer L Bolton, Jennifer L Bragg-Gresham, Steven Buyske, Ayse Demirkan, Guohong Deng, Georg B Ehret, Bjarke Feenstra, Mary F Feitosa, Krista Fischer, Anuj Goel, Jian Gong, Anne U Jackson, Stavroula Kanoni, Marcus E Kleber, Kati Kristiansson, Unhee Lim, Vaneet Lotay, Massimo Mangino, Irene Mateo Leach, Carolina Medina-Gomez, Sarah E Medland, Michael A Nalls, Cameron D Palmer, Dorota Pasko, Sonali Pechlivanis, Marjolein J Peters, Inga Prokopenko, Dmitry Shungin, Alena Stančáková, Rona J Strawbridge, Yun Ju Sung, Toshiko Tanaka, Alexander Teumer, Stella Trompet, Sander W van der Laan, Jessica van Setten, Jana V Van Vliet-Ostaptchouk, Zhaoming Wang, Loïc Yengo, Weihua Zhang, Aaron Isaacs, Eva Albrecht, Johan Ärnlöv, Gillian M Arscott, Antony P Attwood, Stefania Bandinelli, Amy Barrett, Isabelita N Bas, Claire Bellis, Amanda J Bennett, Christian Berne, Roza Blagieva, Matthias Blüher, Stefan Böhringer, Lori L Bonnycastle, Yvonne Böttcher, Heather A Boyd, Marcel Bruinenberg, Ida H Caspersen, Yii-Der Ida Chen, Robert Clarke, E Warwick Daw, Anton J M de Craen, Graciela Delgado, Maria Dimitriou, Alex S F Doney, Niina Eklund, Karol Estrada, Elodie Eury, Lasse Folkersen, Ross M Fraser, Melissa E Garcia, Frank Geller, Vilmantas Giedraitis, Bruna Gigante, Alan S Go, Alain Golay, Alison H Goodall, Scott D Gordon, Mathias Gorski, Hans-Jörgen Grabe, Harald Grallert, Tanja B Grammer, Jürgen Gräßler, Henrik Grönberg, Christopher J Groves, Gaëlle Gusto, Jeffrey Haessler, Per Hall, Toomas Haller, Goran Hallmans, Catharina A Hartman, Maija Hassinen, Caroline Hayward, Nancy L Heard-Costa, Quinta Helmer, Christian Hengstenberg, Oddgeir Holmen, Jouke-Jan Hottenga, Alan L James, Janina M Jeff, Åsa Johansson, Jennifer Jolley, Thorhildur Juliusdottir, Leena Kinnunen, Wolfgang Koenig, Markku Koskenvuo, Wolfgang Kratzer, Jaana Laitinen, Claudia Lamina, Karin Leander, Nanette R Lee, Peter Lichtner, Lars Lind, Jaana Lindström, Ken Sin Lo, Stéphane Lobbens, Roberto Lorbeer, Yingchang Lu, François Mach, Patrik K E Magnusson, Anubha Mahajan, Wendy L McArdle, Stela McLachlan, Cristina Menni, Sigrun Merger, Evelin Mihailov, Lili Milani, Alireza Moayyeri, Keri L Monda, Mario A Morken, Antonella Mulas, Gabriele Müller, Martina Müller-Nurasyid, Arthur W Musk, Ramaiah Nagaraja, Markus M Nöthen, Ilja M Nolte, Stefan Pilz, Nigel W Rayner, Frida Renstrom, Rainer Rettig, Janina S Ried, Stephan Ripke, Neil R Robertson, Lynda M Rose, Serena Sanna, Hubert Scharnagl, Salome Scholtens, Fredrick R Schumacher, William R Scott, Thomas Seufferlein, Jianxin Shi, Albert Vernon Smith, Joanna Smolonska, Alice V Stanton, Valgerdur Steinthorsdottir, Kathleen Stirrups, Heather M Stringham, Johan Sundström, Morris A Swertz, Amy J Swift, Ann-Christine Syvänen, Sian-Tsung Tan, Bamidele O Tayo, Barbara Thorand, Gudmar Thorleifsson, Jonathan P Tyrer, Hae-Won Uh, Liesbeth Vandenput, Frank C Verhulst, Sita H Vermeulen, Niek Verweij, Judith M Vonk, Lindsay L Waite, Helen R Warren, Dawn Waterworth, Michael N Weedon, Lynne R Wilkens, Christina Willenborg, Tom Wilsgaard, Mary K Wojczynski, Andrew Wong, Alan F Wright, Qunyuan Zhang, LifeLines Cohort Study, Eoin P Brennan, Murim Choi, Zari Dastani, Alexander W Drong, Per Eriksson, Anders Franco-Cereceda, Jesper R Gådin, Ali G Gharavi, Michael E Goddard, Robert E Handsaker, Jinyan Huang, Fredrik Karpe, Sekar Kathiresan, Sarah Keildson, Krzysztof Kiryluk, Michiaki Kubo, Jong-Young Lee, Liming Liang, Richard P Lifton, Baoshan Ma, Steven A McCarroll, Amy J McKnight, Josine L Min, Miriam F Moffatt, Grant W Montgomery, Joanne M Murabito, George Nicholson, Dale R Nyholt, Yukinori Okada, John R B Perry, Rajkumar Dorajoo, Eva Reinmaa, Rany M Salem, Niina Sandholm, Robert A Scott, Lisette Stolk, Atsushi Takahashi, Toshihiro Tanaka, Ferdinand M van’t Hooft, Anna A E Vinkhuyzen, Harm-Jan Westra, Wei Zheng, Krina T Zondervan, ADIPOGen Consortium, AGEN-BMI Working Group, CARDIOGRAMplusC4D Consortium, CKDGen Consortium, GLGC, ICBP, MAGIC Investigators, MuTHER Consortium, MIGen Consortium, PAGE Consortium, ReproGen Consortium, GENIE Consortium, International Endogene Consortium, Andrew C Heath, Dominique Arveiler, Stephan J L Bakker, John Beilby, Richard N Bergman, John Blangero, Pascal Bovet, Harry Campbell, Mark J Caulfield, Giancarlo Cesana, Aravinda Chakravarti, Daniel I Chasman, Peter S Chines, Francis S Collins, Dana C Crawford, L Adrienne Cupples, Daniele Cusi, John Danesh, Ulf de Faire, Hester M den Ruijter, Anna F Dominiczak, Raimund Erbel, Jeanette Erdmann, Johan G Eriksson, Martin Farrall, Stephan B Felix, Ele Ferrannini, Jean Ferrières, Ian Ford, Nita G Forouhi, Terrence Forrester, Oscar H Franco, Ron T Gansevoort, Pablo V Gejman, Christian Gieger, Omri Gottesman, Vilmundur Gudnason, Ulf Gyllensten, Alistair S Hall, Tamara B Harris, Andrew T Hattersley, Andrew A Hicks, Lucia A Hindorff, Aroon D Hingorani, Albert Hofman, Georg Homuth, G Kees Hovingh, Steve E Humphries, Steven C Hunt, Elina Hyppönen, Thomas Illig, Kevin B Jacobs, Marjo-Riitta Jarvelin, Karl-Heinz Jöckel, Berit Johansen, Pekka Jousilahti, J Wouter Jukema, Antti M Jula, Jaakko Kaprio, John J P Kastelein, Sirkka M Keinanen-Kiukaanniemi, Lambertus A Kiemeney, Paul Knekt, Jaspal S Kooner, Charles Kooperberg, Peter Kovacs, Aldi T Kraja, Meena Kumari, Johanna Kuusisto, Timo A Lakka, Claudia Langenberg, Loic Le Marchand, Terho Lehtimäki, Valeriya Lyssenko, Satu Männistö, André Marette, Tara C Matise, Colin A McKenzie, Barbara McKnight, Frans L Moll, Andrew D Morris, Andrew P Morris, Jeffrey C Murray, Mari Nelis, Claes Ohlsson, Albertine J Oldehinkel, Ken K Ong, Pamela A F Madden, Gerard Pasterkamp, John F Peden, Annette Peters, Dirkje S Postma, Peter P Pramstaller, Jackie F Price, Lu Qi, Olli T Raitakari, Tuomo Rankinen, D C Rao, Treva K Rice, Paul M Ridker, John D Rioux, Marylyn D Ritchie, Igor Rudan, Veikko Salomaa, Nilesh J Samani, Jouko Saramies, Mark A Sarzynski, Heribert Schunkert, Peter E H Schwarz, Peter Sever, Alan R Shuldiner, Juha Sinisalo, Ronald P Stolk, Konstantin Strauch, Anke Tönjes, David-Alexandre Trégouët, Angelo Tremblay, Elena Tremoli, Jarmo Virtamo, Marie-Claude Vohl, Uwe Völker, Gérard Waeber, Gonneke Willemsen, Jacqueline C Witteman, M Carola Zillikens, Linda S Adair, Philippe Amouyel, Folkert W Asselbergs, Themistocles L Assimes, Murielle Bochud, Bernhard O Boehm, Eric Boerwinkle, Stefan R Bornstein, Erwin P Bottinger, Claude Bouchard, Stéphane Cauchi, John C Chambers, Stephen J Chanock, Richard S Cooper, Paul I W de Bakker, George Dedoussis, Luigi Ferrucci, Paul W Franks, Philippe Froguel, Leif C Groop, Christopher A Haiman, Anders Hamsten, Jennie Hui, David J Hunter, Kristian Hveem, Robert C Kaplan, Mika Kivimaki, Diana Kuh, Markku Laakso, Yongmei Liu, Nicholas G Martin, Winfried März, Mads Melbye, Andres Metspalu, Susanne Moebus, Patricia B Munroe, Inger Njølstad, Ben A Oostra, Colin N A Palmer, Nancy L Pedersen, Markus Perola, Louis Pérusse, Ulrike Peters, Chris Power, Thomas Quertermous, Rainer Rauramaa, Fernando Rivadeneira, Timo E Saaristo, Danish Saleheen, Naveed Sattar, Eric E Schadt, David Schlessinger, P Eline Slagboom, Harold Snieder, Tim D Spector, Unnur Thorsteinsdottir, Michael Stumvoll, Jaakko Tuomilehto, André G Uitterlinden, Matti Uusitupa, Pim van der Harst, Mark Walker, Henri Wallaschofski, Nicholas J Wareham, Hugh Watkins, David R Weir, H-Erich Wichmann, James F Wilson, Pieter Zanen, Ingrid B Borecki, Panos Deloukas, Caroline S Fox, Iris M Heid, Jeffrey R O’Connell, David P Strachan, Kari Stefansson, Cornelia M van Duijn, Gonçalo R Abecasis, Lude Franke, Timothy M Frayling, Mark I McCarthy, Peter M Visscher, André Scherag, Cristen J Willer, Michael Boehnke, Karen L Mohlke, Cecilia M Lindgren, Jacques S Beckmann, Inês Barroso, Kari E North, Erik Ingelsson, Joel N Hirschhorn, Ruth J F Loos, and Elizabeth K Speliotes. Genetic studies of body mass index yield new insights for obesity biology. Nature, 518(7538):197–206, February 2015.

[31] Loic Yengo, Julia Sidorenko, Kathryn E Kemper, Zhili Zheng, Andrew R Wood, Michael N Weedon, Timothy M Frayling, Joel Hirschhorn, Jian Yang, Peter M Visscher, and the GIANT Consortium. Meta-analysis of genome-wide association studies for height and body mass index in ~700 000 individuals of european ancestry. Human Molecular Genetics, 2018.

[32] Karen A Schlauch, Robert W Read, Vincent C Lombardi, Gai Elhanan, William J Metcalf, Anthony D Slonim, 23andMe Research Team, and Joseph J Grzymski. A comprehensive Genome-Wide and Phenome-Wide examination of BMI and obesity in a northern nevadan cohort. G3, 10(2):645–664, February 2020.

[33] S Miyata, N Matsumoto, K Taguchi, A Akagi, T Iino, N Funatsu, and S Maekawa. Biochemical and ultrastructural analyses of IgLON cell adhesion molecules, kilon and OBCAM in the rat brain. Neuroscience, 117(3):645–658, 2003.

[34] Takashi Hashimoto, Mayumi Yamada, Shohei Maekawa, Toshihiro Nakashima, and Seiji Miyata. IgLON cell adhesion molecule kilon is a crucial modulator for synapse number in hippocampal neurons. Brain Res., 1224:1–11, August 2008.

[35] Ricardo Sanz, Gino B Ferraro, and Alyson E Fournier. IgLON cell adhesion molecules are shed from the cell surface of cortical neurons to promote neuronal growth. J. Biol. Chem., 290(7):4330–4342, February 2015.

[36] Angela W S Lee, Heidi Hengstler, Kathrin Schwald, Mauricio Berriel-Diaz, Desirée Loreth, Matthias Kirsch, Oliver Kretz, Carola A Haas, Martin Hrabě de Angelis, Stephan Herzig, Thomas Brümmendorf, Martin Klingenspor, Fritz G Rathjen, Jan Rozman, George Nicholson, Roger D Cox, and Michael K E Schäfer. Functional inactivation of the genome-wide association study obesity gene neuronal growth regulator 1 in mice causes a body mass phenotype. PLoS One, 7(7):e41537, July 2012.

[37] Katyayani Singh, Desirée Loreth, Bruno Pöttker, Kyra Hefti, Jürgen Innos, Kathrin Schwald, Heidi Hengstler, Lutz Menzel, Clemens J Sommer, Konstantin Radyushkin, Oliver Kretz, Mari-Anne Philips, Carola A Haas, Katrin Frauenknecht, Kersti Lilleväli, Bernd Heimrich, Eero Vasar, and Michael K E Schäfer. Neuronal growth and behavioral alterations in mice deficient for the psychiatric Disease-Associated negr1 gene. Front. Mol. Neurosci., 11:30, February 2018.

[38] Elizabeth K Speliotes, Cristen J Willer, Sonja I Berndt, Keri L Monda, Gudmar Thorleifsson, Anne U Jackson, Hana Lango Allen, Cecilia M Lindgren, Jian’an Luan, Reedik Mägi, Joshua C Randall, Sailaja Vedantam, Thomas W Winkler, Lu Qi, Tsegaselassie Workalemahu, Iris M Heid, Valgerdur Steinthorsdottir, Heather M Stringham, Michael N Weedon, Eleanor Wheeler, Andrew R Wood, Teresa Ferreira, Robert J Weyant, Ayellet V Segrè, Karol Estrada, Liming Liang, James Nemesh, Ju-Hyun Park, Stefan Gustafsson, Tuomas O Kilpeläinen, Jian Yang, Nabila Bouatia-Naji, Tõnu Esko, Mary F Feitosa, Zoltán Kutalik, Massimo Mangino, Soumya Raychaudhuri, Andre Scherag, Albert Vernon Smith, Ryan Welch, Jing Hua Zhao, Katja K Aben, Devin M Absher, Najaf Amin, Anna L Dixon, Eva Fisher, Nicole L Glazer, Michael E Goddard, Nancy L Heard-Costa, Volker Hoesel, Jouke-Jan Hottenga, Asa Johansson, Toby Johnson, Shamika Ketkar, Claudia Lamina, Shengxu Li, Miriam F Moffatt, Richard H Myers, Narisu Narisu, John R B Perry, Marjolein J Peters, Michael Preuss, Samuli Ripatti, Fernando Rivadeneira, Camilla Sandholt, Laura J Scott, Nicholas J Timpson, Jonathan P Tyrer, Sophie van Wingerden, Richard M Watanabe, Charles C White, Fredrik Wiklund, Christina Barlassina, Daniel I Chasman, Matthew N Cooper, John-Olov Jansson, Robert W Lawrence, Niina Pellikka, Inga Prokopenko, Jianxin Shi, Elisabeth Thiering, Helene Alavere, Maria T S Alibrandi, Peter Almgren, Alice M Arnold, Thor Aspelund, Larry D Atwood, Beverley Balkau, Anthony J Balmforth, Amanda J Bennett, Yoav Ben-Shlomo, Richard N Bergman, Sven Bergmann, Heike Biebermann, Alexandra I F Blakemore, Tanja Boes, Lori L Bonnycastle, Stefan R Bornstein, Morris J Brown, Thomas A Buchanan, Fabio Busonero, Harry Campbell, Francesco P Cappuccio, Christine Cavalcanti-Proença, Yii-Der Ida Chen, Chih-Mei Chen, Peter S Chines, Robert Clarke, Lachlan Coin, John Connell, Ian N M Day, Martin den Heijer, Jubao Duan, Shah Ebrahim, Paul Elliott, Roberto Elosua, Gudny Eiriksdottir, Michael R Erdos, Johan G Eriksson, Maurizio F Facheris, Stephan B Felix, Pamela Fischer-Posovszky, Aaron R Folsom, Nele Friedrich, Nelson B Freimer, Mao Fu, Stefan Gaget, Pablo V Gejman, Eco J C Geus, Christian Gieger, Anette P Gjesing, Anuj Goel, Philippe Goyette, Harald Grallert, Jürgen Grässler, Danielle M Greenawalt, Christopher J Groves, Vilmundur Gudnason, Candace Guiducci, Anna-Liisa Hartikainen, Neelam Hassanali, Alistair S Hall, Aki S Havulinna, Caroline Hayward, Andrew C Heath, Christian Hengstenberg, Andrew A Hicks, Anke Hinney, Albert Hofman, Georg Homuth, Jennie Hui, Wilmar Igl, Carlos Iribarren, Bo Isomaa, Kevin B Jacobs, Ivonne Jarick, Elizabeth Jewell, Ulrich John, Torben Jørgensen, Pekka Jousilahti, Antti Jula, Marika Kaakinen, Eero Kajantie, Lee M Kaplan, Sekar Kathiresan, Johannes Kettunen, Leena Kinnunen, Joshua W Knowles, Ivana Kolcic, Inke R König, Seppo Koskinen, Peter Kovacs, Johanna Kuusisto, Peter Kraft, Kirsti Kvaløy, Jaana Laitinen, Olivier Lantieri, Chiara Lanzani, Lenore J Launer, Cecile Lecoeur, Terho Lehtimäki, Guillaume Lettre, Jianjun Liu, Marja-Liisa Lokki, Mattias Lorentzon, Robert N Luben, Barbara Ludwig, MAGIC, Paolo Manunta, Diana Marek, Michel Marre, Nicholas G Martin, Wendy L McArdle, Anne McCarthy, Barbara McKnight, Thomas Meitinger, Olle Melander, David Meyre, Kristian Midthjell, Grant W Montgomery, Mario A Morken, Andrew P Morris, Rosanda Mulic, Julius S Ngwa, Mari Nelis, Matt J Neville, Dale R Nyholt, Christopher J O’Donnell, Stephen O’Rahilly, Ken K Ong, Ben Oostra, Guillaume Paré, Alex N Parker, Markus Perola, Irene Pichler, Kirsi H Pietiläinen, Carl G P Platou, Ozren Polasek, Anneli Pouta, Suzanne Rafelt, Olli Raitakari, Nigel W Rayner, Martin Ridderstråle, Winfried Rief, Aimo Ruokonen, Neil R Robertson, Peter Rzehak, Veikko Salomaa, Alan R Sanders, Manjinder S Sandhu, Serena Sanna, Jouko Saramies, Markku J Savolainen, Susann Scherag, Sabine Schipf, Stefan Schreiber, Heribert Schunkert, Kaisa Silander, Juha Sinisalo, David S Siscovick, Jan H Smit, Nicole Soranzo, Ulla Sovio, Jonathan Stephens, Ida Surakka, Amy J Swift, Mari-Liis Tammesoo, Jean-Claude Tardif, Maris Teder-Laving, Tanya M Teslovich, John R Thompson, Brian Thomson, Anke Tönjes, Tiinamaija Tuomi, Joyce B J van Meurs, Gert-Jan van Ommen, Vincent Vatin, Jorma Viikari, Sophie Visvikis-Siest, Veronique Vitart, Carla I G Vogel, Benjamin F Voight, Lindsay L Waite, Henri Wallaschofski, G Bragi Walters, Elisabeth Widen, Susanna Wiegand, Sarah H Wild, Gonneke Willemsen, Daniel R Witte, Jacqueline C Witteman, Jianfeng Xu, Qunyuan Zhang, Lina Zgaga, Andreas Ziegler, Paavo Zitting, John P Beilby, I Sadaf Farooqi, Johannes Hebebrand, Heikki V Huikuri, Alan L James, Mika Kähönen, Douglas F Levinson, Fabio Macciardi, Markku S Nieminen, Claes Ohlsson, Lyle J Palmer, Paul M Ridker, Michael Stumvoll, Jacques S Beckmann, Heiner Boeing, Eric Boerwinkle, Dorret I Boomsma, Mark J Caulfield, Stephen J Chanock, Francis S Collins, L Adrienne Cupples, George Davey Smith, Jeanette Erdmann, Philippe Froguel, Henrik Grönberg, Ulf Gyllensten, Per Hall, Torben Hansen, Tamara B Harris, Andrew T Hattersley, Richard B Hayes, Joachim Heinrich, Frank B Hu, Kristian Hveem, Thomas Illig, Marjo-Riitta Jarvelin, Jaakko Kaprio, Fredrik Karpe, Kay-Tee Khaw, Lambertus A Kiemeney, Heiko Krude, Markku Laakso, Debbie A Lawlor, Andres Metspalu, Patricia B Munroe, Willem H Ouwehand, Oluf Pedersen, Brenda W Penninx, Annette Peters, Peter P Pramstaller, Thomas Quertermous, Thomas Reinehr, Aila Rissanen, Igor Rudan, Nilesh J Samani, Peter E H Schwarz, Alan R Shuldiner, Timothy D Spector, Jaakko Tuomilehto, Manuela Uda, André Uitterlinden, Timo T Valle, Martin Wabitsch, Gérard Waeber, Nicholas J Wareham, Hugh Watkins, Procardis Consortium, James F Wilson, Alan F Wright, M Carola Zillikens, Nilanjan Chatterjee, Steven A McCarroll, Shaun Purcell, Eric E Schadt, Peter M Visscher, Themistocles L Assimes, Ingrid B Borecki, Panos Deloukas, Caroline S Fox, Leif C Groop, Talin Haritunians, David J Hunter, Robert C Kaplan, Karen L Mohlke, Jeffrey R O’Connell, Leena Peltonen, David Schlessinger, David P Strachan, Cornelia M van Duijn, H-Erich Wichmann, Timothy M Frayling, Unnur Thorsteinsdottir, Gonçalo R Abecasis, Inês Barroso, Michael Boehnke, Kari Stefansson, Kari E North, Mark I McCarthy, Joel N Hirschhorn, Erik Ingelsson, and Ruth J F Loos. Association analyses of 249,796 individuals reveal 18 new loci associated with body mass index. Nat. Genet., 42(11):937–948, November 2010.

[39] Craig L Hyde, Michael W Nagle, Chao Tian, Xing Chen, Sara A Paciga, Jens R Wendland, Joyce Y Tung, David A Hinds, Roy H Perlis, and Ashley R Winslow. Identification of 15 genetic loci associated with risk of major depression in individuals of european descent. Nat. Genet., 48(9):1031–1036, September 2016.

[40] D M Sabatini, R K Barrow, S Blackshaw, P E Burnett, M M Lai, M E Field, B A Bahr, J Kirsch, H Betz, and S H Snyder. Interaction of RAFT1 with gephyrin required for rapamycin-sensitive signaling. Science, 284(5417):1161–1164, May 1999.

[41] Franz Vauti, Tobias Goller, Rafael Beine, Lore Becker, Thomas Klopstock, Sabine M Hölter, Wolfgang Wurst, Helmut Fuchs, Valerie Gailus-Durner, Martin Hrabé de Angelis, and Hans-Henning Arnold. The mouse trm1-like gene is expressed in neural tissues and plays a role in motor coordination and exploratory behaviour. Gene, 389(2):174–185, March 2007.

[42] Francisca Salas-Pérez, Omar Ramos-Lopez, María L Mansego, Fermín I Milagro, José L Santos, José I Riezu-Boj, and J Alfredo Martínez. DNA methylation in genes of longevity-regulating pathways: association with obesity and metabolic complications. Aging, 11(6):1874–1899, March 2019.

[43] Laura Braccini, Elisa Ciraolo, Carlo C Campa, Alessia Perino, Dario L Longo, Gianpaolo Tibolla, Marco Pregnolato, Yanyan Cao, Beatrice Tassone, Federico Damilano, Muriel Laffargue, Enzo Calautti, Marco Falasca, Giuseppe D Norata, Jonathan M Backer, and Emilio Hirsch. PI3K-C2*γ* is a rab5 effector selectively controlling endosomal akt2 activation downstream of insulin signalling. Nat. Commun., 6:7400, June 2015.

[44] Nadine N Hauer, Bernt Popp, Leila Taher, Carina Vogl, Perundurai S Dhandapany, Christian Büttner, Steffen Uebe, Heinrich Sticht, Fulvia Ferrazzi, Arif B Ekici, Alessandro De Luca, Patrizia Klinger, Cornelia Kraus, Christiane Zweier, Antje Wiesener, Rami Abou Jamra, Erdmute Kunstmann, Anita Rauch, Dagmar Wieczorek, Anna-Marie Jung, Tilman R Rohrer, Martin Zenker, Helmuth-Guenther Doerr, André Reis, and Christian T Thiel. Evolutionary conserved networks of human height identify multiple mendelian causes of short stature. Eur. J. Hum. Genet., 27(7):1061–1071, July 2019.

[45] Masahiro Horita, Keiichiro Nishida, Joe Hasei, Takayuki Furumatsu, Miwa Sakurai, Yuta Onodera, Kanji Fukuda, Donald M Salter, and Toshifumi Ozaki. Involvement of ADAM12 in chondrocyte differentiation by regulation of TGF-*β*1-Induced IGF-1 and RUNX-2 expressions. Calcif. Tissue Int., 105(1):97–106, July 2019.

[46] Amelie Baud, Megan K Mulligan, Francesco Paolo Casale, Jesse F Ingels, Casey J Bohl, Jacques Callebert, Jean-Marie Launay, Jon Krohn, Andres Legarra, Robert W Williams, and Oliver Stegle. Genetic variation in the social environment contributes to health and disease. PLoS Genet., 13(1):e1006498, January 2017.

[47] Chen-Yu Liao, Brad A Rikke, Thomas E Johnson, Jonathan A L Gelfond, Vivian Diaz, and James F Nelson. Fat maintenance is a predictor of the murine lifespan response to dietary restriction. Aging Cell, 10(4):629–639, August 2011.

